# Discovering cryptic pocket opening and binding of a stimulant derivative in a vestibular site of the 5-HT_3_*_A_* receptor

**DOI:** 10.1101/2023.11.13.566806

**Authors:** Nandan Haloi, Emelia Karlsson, Marc Delarue, Rebecca J. Howard, Erik Lindahl

## Abstract

Ligand-gated ion channels propagate electrochemical signals in the nervous system. A diverse set of modulators including stimulants, anesthetics, and lipids regulate their function; however, structures of ligand-bound complexes can be difficult to capture by experimental methods, particularly when binding is dynamic or transient. Here, we used computational methods and electrophysiology to identify a possible bound state of a modulatory stimulant derivative in a cryptic vestibular pocket, distinct from the orthosteric neurotransmitter binding site, of a mammalian serotonin-3A receptor. Starting from a closed-pocket experimental structure, we first applied a molecular dynamics simulations-based goal-oriented adaptive sampling method to identify possible open-pocket conformations. To find plausible ligand-binding poses, we performed Boltzmann docking, which combines traditional docking with Markov state modeling, of the newly sampled conformations. Clustering and analysis of stability and accessibility of docked poses supported a preferred binding site; we further validated this site by mutagenesis and electrophysiology, suggesting a mechanism of potentiation by stabilizing intersubunit contacts. Given the pharmaceutical relevance of serotonin-3 receptors in emesis, psychiatric and gastrointestinal diseases, characterizing relatively unexplored modulatory sites such as these could open valuable avenues to understanding conformational cycling and designing state-dependent drugs.

**Significance:** 5-HT_3A_ receptors receive the chemical signals of excitatory neurotransmission across the synapse in the central and peripheral nervous systems, and are involved in conditions including emesis, pain, psychiatric disorders, drug abuse, and irritable bowel syndrome. Given their pharmaceutical importance, there is great interest in understanding how and where ligands interact with these receptors. A pocket facing the extracellular vestibule of this membrane protein has been proposed as a modulatory site, but it remains largely uncharacterized in the context of structural modeling or pharmacologically relevant ligands. Here, we are able to identify and investigate binding of a stimulant derivative, 4-bromoamphetamine, in this site by using an integrative computational and experimental approach that is able to account for conformational flexibility.

## Introduction

Pentameric ligand-gated ion channels (pLGICs) play central roles in intercellular communication in the mammalian nervous system [1]. In a classical example, neurotransmitters released at the synaptic cleft bind to corresponding pLGICs, contracting the extracellular domain (ECD) and opening a permeation pore in the transmembrane domain (TMD) to allow ions to cross the membrane for further signal transduction [2] (Fig. 1A). A diverse set of ligands, including anesthetics, neurosteroids, and lipids, can modulate these proteins via orthosteric (Fig. 1A, yellow) or various allosteric modulatory sites. For example, in serotonin-3 receptors (5-HT_3_Rs), antiemetic drugs such as palonosetron, used in the treatment of nausea and vomiting associated with radiation and chemotherapies, compete with serotonin binding at the orthosteric site to inhibit channel function [3]. Conversely, in type-A *γ*-aminobutyric acid receptors (GABA_A_Rs), classical benzodiazepine drugs that are used to treat epilepsy, anxiety, and insomnia bind sites distinct from that of GABA to modulate activation [4–6].

**Fig. 1.**
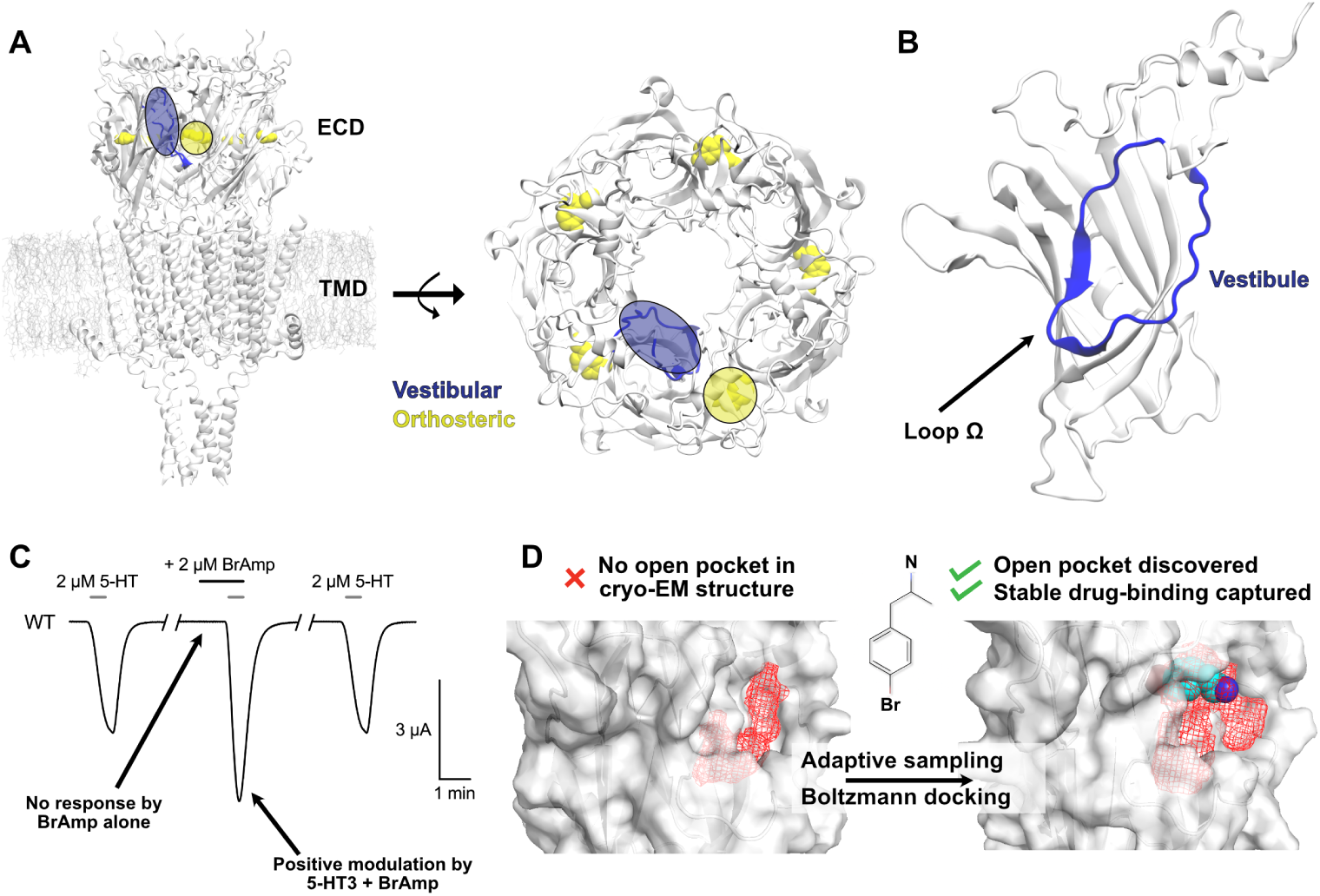
Overview of 5-HT_3A_R structure and pharmacology explored in this work. A) Architecture of a representative 5-HT_3A_R, colored by subunit, viewed from the membrane plane (left) and from the extracellular side (right). Serotonin bound at the orthosteric site is represented in yellow. The vestibular site is colored blue. B) ECD of a single 5-HT_3A_R subunit, with the Ω-loop in blue. C) Representative current trace from a 5-HT_3A_R-expressing *Xenopus* oocyte in the absence and presence of 4-bromoamphetamine (BrAmp) during serotonin (at EC_20_) pulses. D) Pocket volumes at the 5-HT_3A_R vestibular site, generated in Fpocket [23], show a superficial cavity in the experimental structure (*left*), but a clearly opened pocket surrounding the docked pose of 4-bromoamphetamine (*right*), as found in our study. Pocket volumes are shown in red mesh, the ligand in van der Waals, and the receptor in gray surface representations.

Whereas both the orthosteric neurotransmitter site and ECD benzodiazepine site are located at extracellular subunit interfaces, the ECD interior vestibule constitutes a relatively unexplored potential modulatory region (Fig. 1A, blue). Despite the recent determination of several ligand-bound experimental structures, none have yet captured binding at the vestibular site in eukaryotic pLGICs. In contrast, several X-ray structures of prokaryotic pLGICs do resolve ligands in this region, including the positive modulators 4-bromocinnamate (PDB ID: 6FLI) and 4-bromoamphetamine (PDB ID: 9EWL) bound to sTeLIC, a channel derived from an endosymbiont of the tubeworm *Tevnia jerichonana* [7, 8] (Fig. 1C). Similarly, in the plant-pathogen channel ELIC and proton-gated channel GLIC, the benzodiazepine flurazepam (PDB ID: 2YOE, chain C) [9] and various modulatory carboxylates (PDB IDs: 6HJZ, 4QH1, 6HPP), respectively, were found to bind in this location (Fig. S1) [10–12]. In these cases, ligands were identified between the *β*4, *β*5, and *β*6 strands, in a vestibular cavity defined by the so-called Ω-loop (Fig. 1B).

In an effort to investigate a role for the ECD vestibular site in ligand binding and modulation of eukaryotic pLGICs, Brams and colleagues [13] performed a systematic analysis using available experimental structures. As demonstrated in their study, this cavity is occluded in experimental structures of most eukaryotic pLGICs, including *α*2/3 nicotinic acetylcholine receptor subunits [14, 15] and *α*1/3 glycine receptor subunits [16]. Interestingly, the vestibular cavity appears to be more accessible in 5-HT_3A_Rs [17]; indeed, cysteine-scanning mutagenesis of human 5-HT_3A_Rs indicated this region to be sensitive to modulation by covalent labeling [13]. However, the 5-HT_3A_R Ω-loop is not conserved with that of prokaryotic channels known to bind vestibular ligands (Fig. S1), the pocket conformation present in the apo structure might not be compatible with modulator binding, and direct evidence for drug modulation via this region in a eukaryotic system remains lacking.

Molecular dynamics (MD)-based techniques offer complementary approaches to explore binding sites not readily apparent in experimental structures [18–21]. Here, we applied a goal-oriented adaptive sampling method [22] to explore regions of conformational space relevant to the potential opening of this pocket in the ECD vestibule of a mammalian 5-HT_3A_R (Fig. 1C-D). Then, to find plausible ligand-binding poses, we performed ensemble docking of 4-bromoamphetamine resulting in a total of 1.5 million docked poses, and reweighted docking scores by Boltzmann-distribution probabilities of each protein conformation occurring, as derived from Markov state model analysis of our trajectories. Following clustering of the top 100 docked poses, we then performed replicate unbiased MD simulations of representative complexes in two forcefields to estimate ligand stability, and screened the most stable complexes for pocket accessibility to the aqueous environment. For one relatively stable and accessible site, mutations predicted to disrupt 4-bromoamphetamine binding and/or modulation were validated by electrophysiology recordings in *Xenopus laevis* oocytes, and provided a mechanistic rationale for the modulatory stabilization of an activated state.

## Results

### Evidence for 5-HT3AR potentiation from functional but not structural data

In exploring potential vestibular pLGIC modulators, we noted that ligands previously crystallized in the vestibular site of the prokaryotic channel sTeLIC, i.e. 4-bromocinnamate, 4-bromophenethylamine, and 4-bromoamphetamine [7, 8], are structurally similar to amphetamine psychostimulants. We tested the effect of 4-bromoamphetamine on 5-HT_3A_Rs expressed in *Xenopus* oocytes, and found that low micromolar concentrations of the drug enhanced submaximal (2 *µ*M, EC20) 5-HT currents (Fig. 1C), with even greater apparent potency than in the bacterial channel [8]. In the absence of 5-HT, 4-bromoamphetamine did not directly activate the receptor at up to 200 *µ*M, suggesting it acts not at the orthosteric 5-HT site but rather via an allosteric modulatory site (Fig. 1C, Fig. S2). However, we found that the the vestibular site shown to bind 4-bromoamphetamine in sTeLIC was relatively contracted in experimental 5-HT_3A_R structures, even in an activated state presumed to be stabilized by positive allosteric modulators (PDB IDs: 6DG8 and 4PIR) (Fig. 1D, Fig. S3) [8, 13, 24]. Solvent-accessible volume in this pocket was limited; although smaller dicarboxylates could fit (based on structural alignment with GLIC structures), ligands on the scale of 4-bromoamphetamine or larger (4-bromocinnamate, flurazepam) superimposed from complexes with sTeLIC and ELIC, respectively, clashed with 5-HT_3A_R Ω-loop sidechains Fig. S1).

To more thoroughly investigate potential binding in this site, we performed molecular docking around the Ω-loop of an activated 5-HT_3A_R structure, and subjected this complex to unrestrained MD simulations. The modulator deviated rapidly (within 20 ns) and dramatically (*>*15 ^°^A root-mean-square deviation, RMSD) from the docked pose, and sampled increasingly distant poses over time (Fig. S4). If the 5-HT_3A_R vestibular site does mediate 4-bromoamphetamine modulation, it would appear to involve a cryptic pocket, i.e. a binding site that is intrinsically transient or otherwise not readily apparent in experimental structures [18]. We also ran 25 replicate unrestrained simulations in the absence of modulator, totaling 1 *µ*s, but observed limited flexibility in the Ω-loop (root-mean-square fluctuation *<* 1.8 ^°^A, Fig. S5) relative to other regions, indicating the conformational transitions are slow and that enhanced sampling methods would be required to capture an alternative open state of this putative binding site.

### An adaptive sampling and docking-based workflow to capture a stable, accessible ligand site

We next applied an adaptive sampling method, fluctuation amplification of specific traits (FAST) [22], to launch simulations from the closed-pocket 5-HT_3A_R structure and maximize the chance of discovering cryptic pockets that may harbor new ligand binding sites (see Methods for details) (Fig. 2A). Within 30 simulation generations, corresponding to 30 *µ*s of total simulation, we observed an increase in pairwise distances between residues in the Ω-loop and neighboring *β*-strand walls, associated with a widening of the Ω-loop as well as more modest increases in pocket volume and solvent-accessible surface area (Fig. 2A, Fig. S6).

**Fig. 2.**
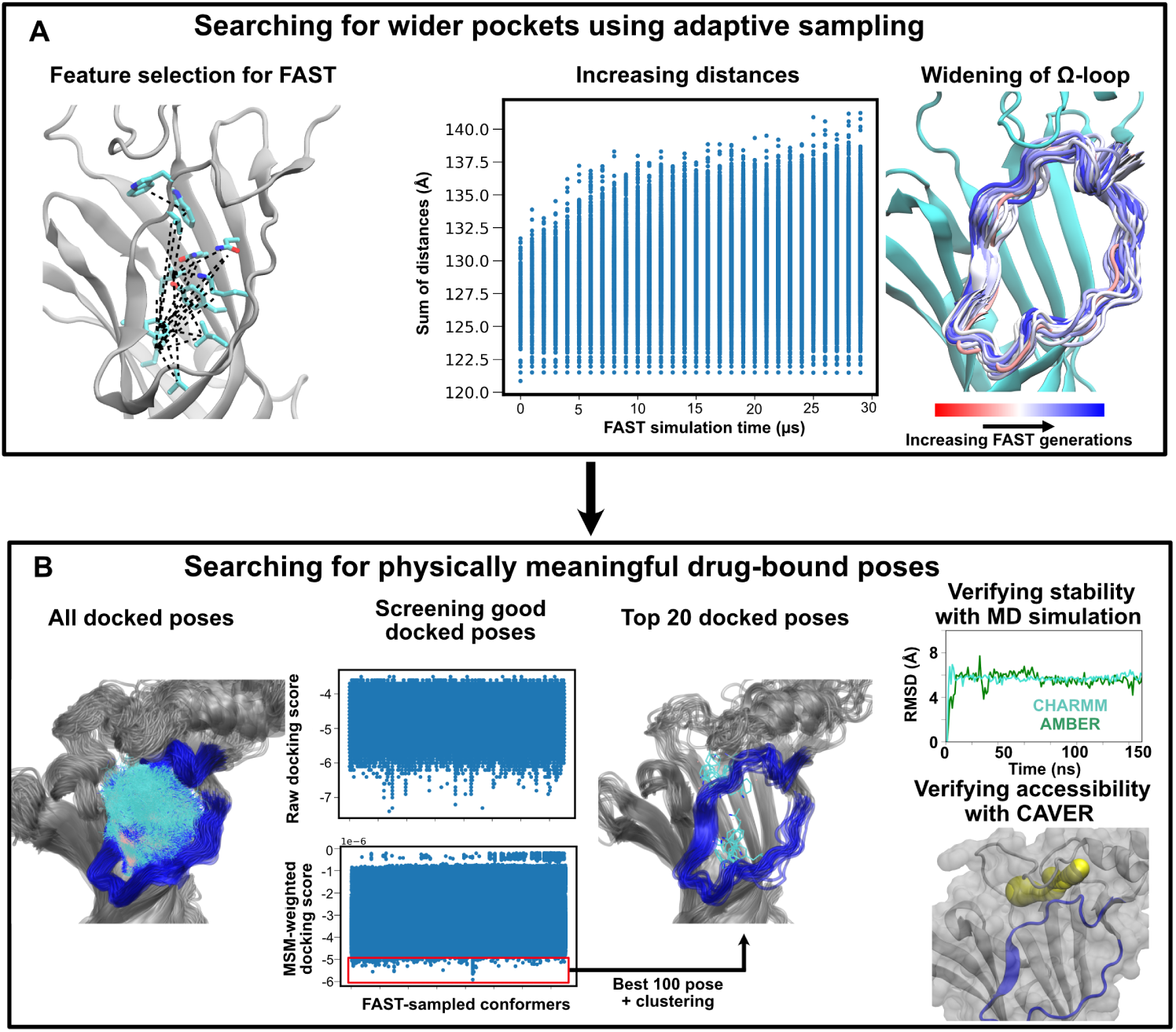
A two-phase computational workflow developed in this study to model first pocket opening and then ligand binding. A) Left: in FAST sampling, starting points for successive generations were selected on the basis of pairwise distances (dashed lines) between residues (cyan) in the Ω-loop and neighboring *β*-strands. Center: these distances increased in successive FAST generations, each containing 1 *µ*s of MD simulations data. Right: FAST sampling was also associated with progressive widening of the Ω-loop, as depicted by representative snapshots colored by generation (red to blue). B) Left: FAST-generated conformers were subjected to molecular docking with 4-bromoamphetamine. Center: docked scores were then re-weighted by the stationary probability of the corresponding protein conformation, determined by Markov state model analysis. The 100 best poses (red box) were then clustered based on RMSD of the ligand atoms with a cutoff of 2.5 °A, and screened down to 20 representative poses. Right: these poses were then used for further MD simulations and accessibility analysis.

To assess ligand binding to these newly explored conformations, we next performed ensemble docking of 4-bromoamphetamine to all FAST-sampled MD trajectories, a total of 30 *µ*s simulations (Fig. 2B). Although a docking score can indicate the likelihood of binding a particular pocket conformation, its contribution to the actual free energy of binding will be low if this pocket conformation only has a low probability of occurring. Hence, we re-weighted the docking scores according to the equilibrium probability of the corresponding protein conformations based on a Markov state model of the FAST-sampled trajectories (Fig. 2B). This approach, termed Boltzmann docking, has been shown to improve activity predictions for small molecules in targets such as TEM *β*-lactamase [25].

From the 100 best Boltzmann-docked 4-bromoamphetamine complexes, we reduced redundancy using RMSD-based clustering, resulting in twenty representative putative poses (Fig. 2B). We then tested their stability by performing unbiased MD simulations. Since small-molecule parameters can be sensitive to the choice of forcefields, we performed three replicate simulations of each system in both CHARMM36m [26–29] and AMBER [30, 31], for a total of 20 systems x 3 replicates x 2 forcefields = 120 150-ns trajectories (Fig. S8).

For closer analysis of potential bound poses, we selected systems in which the ligand remained relatively stable (*<*15 ^°^A center-of-mass RMSD) throughout at least 5 of its 6 replicate simulations. In the six systems that passed this filtering (Fig. S8), we further analyzed the contact frequency of the ligand with each receptor residue. Interestingly, one of these systems (pose 20) positioned the ligand between the *β*2, *β*4 and *β*6 strands (site 1, Fig. 3A–B). In the remaining five relatively stable systems (poses 8, 12, 13, 16, 18), the ligand occupied a distinct region (site 2, Fig. 3B–C) defined by the *β*1, *β*2, *β*4, and *β*8 strands. Notably, the RMSD of the ligand in site 1 was moderately higher (around 5 ^°^A, Fig. 3B) since the ligand changed orientation slightly in the simulation compared to the initial docked pose, converging to the end pose seen in Fig. 3A.

**Fig. 3.**
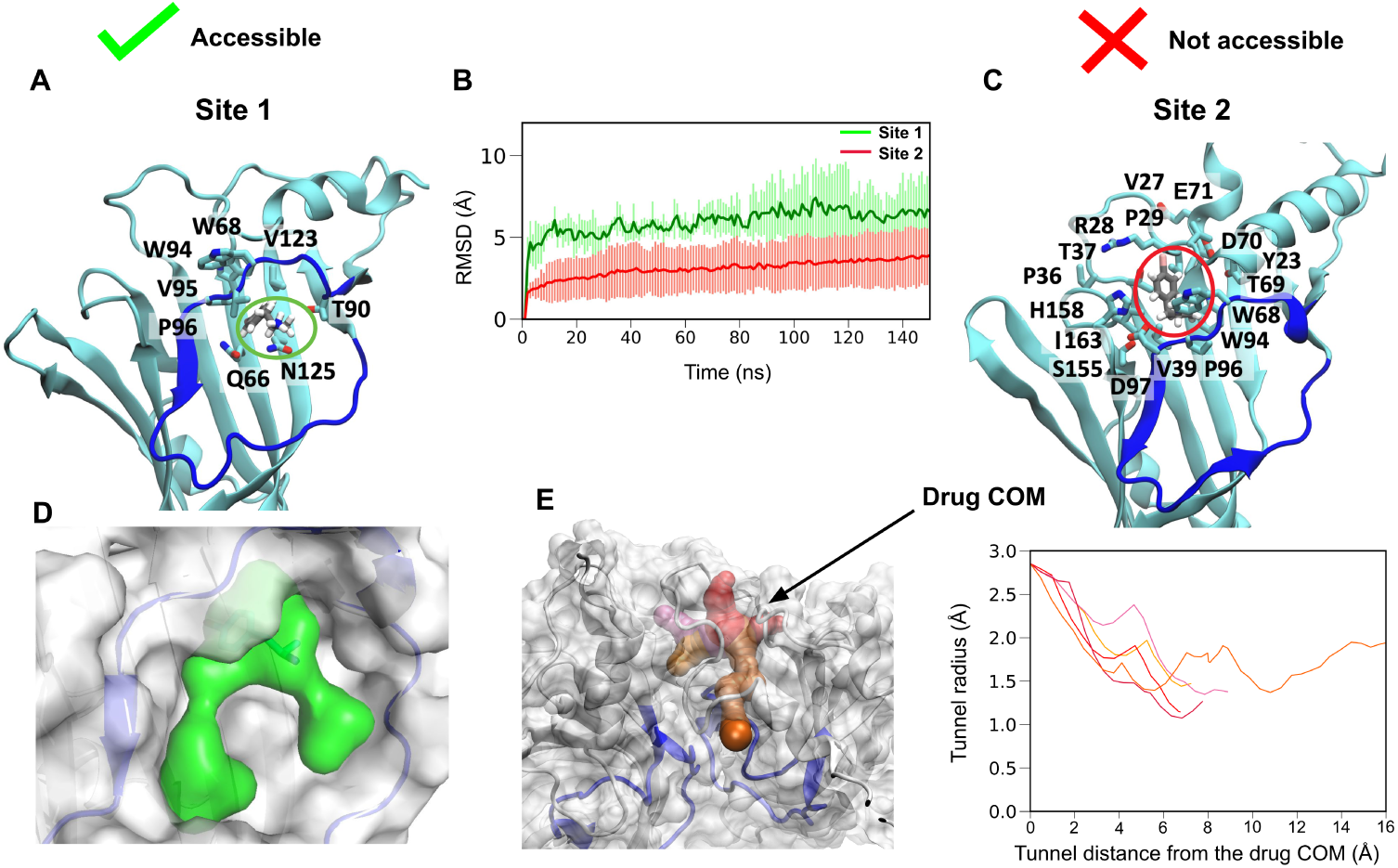
Comparison of stable sites captured by MD simulations of Boltzmann docked poses. A) Interacting residues (cyan, *>*50 % contact probability in MD simulations) surrounding the ligand (gray) in site 1. For clarity, hydrogens are hidden in protein side chains, and the Ω-loop backbone is shown in blue. B) Ligand stability in sites 1 (green) and 2 (red), as represented by RMSD of 4-bromoamphetamine from its initial docked pose during MD simulations, averaged across all stable replicates. Shaded regions indicate standard deviations. C) Interacting residues surrounding the ligand in site 2, depicted as in *A*. This site is located peripheral to the main vestibular pocket. D) Based on accessibility analysis in CAVER [32], the ligand pocket (green) in site 1 is clearly accessible to the solvent. E) In site 2, molecular representations (left) and radius plots (right) show no open accessible pathways (shades of red) to the ligand center of mass.

Although even more stable than site 1, poses in site 2 were buried in an enclosed cavity, substantially shielded from solvent by the outermost ECD (Fig. 3B-C). We noted that computational docking and simulations might retain a ligand in a pocket entirely inaccessible from the solvent, although this makes actual exchange and binding physically implausible. Therefore, we also analyzed the accessibility of both putative sites using the CAVER software suit [32] (Fig. 2B). In site 1, the ligand amino group was directly exposed to solvent, with the pocket clearly accessible to the aqueous medium (Fig. 3D). However, in site 2, accessibility analysis showed no pathways for ligand entry (Fig. 3E). Accordingly, we proceeded to focus on site 1 as the physically most plausible region for ligand binding and modulation.

### Structural features and functional validation of a vestibular modulatory site

The putative 5-HT_3A_R vestibular site contained several hydrophobic residues (*β*2-W68, *β*4-W94, *β*6-V123) lining the inner wall, making van der Waals contacts with the bromine and aromatic groups of the ligand (Fig. 3A–B). The mouth of the putative site contained more polar residues (*β*2-Q66, *β*3-T90, *β*6-N125), interacting with the ligand amino group. This binding pose was comparable to that observed in X-ray structure of this and related ligands in sTeLIC (Fig. 4A,B). In comparing the experimental structure to the putative bound state derived from our simulations, the most dramatic change in the Ω-loop was an outward shift and rotation of the *β*4 strand, relieving prospective clashes of the ligand with residues including *β*4-V95 (Fig. 1C, Fig. 4A).

**Fig. 4.**
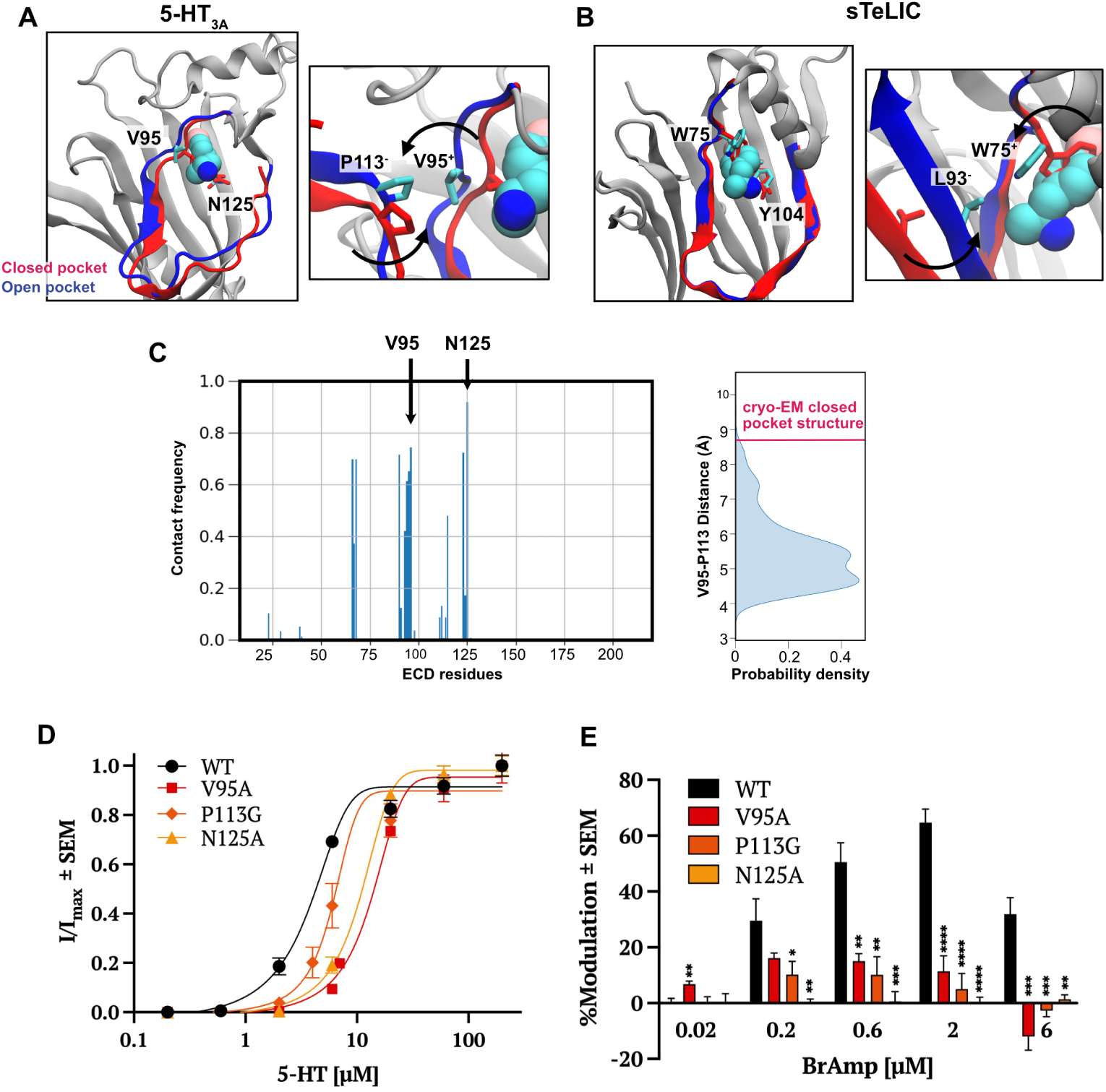
Structural features and functional validation of a vestibular modulatory site of 5-HT_3A_R. A, *left*) Remodeling of the *β*4-strand in open- (blue) versus closed-pocket (red) models of 5-HT_3A_R enables stable binding of the 4-bromoamphetamine (cyan spheres) between residues V95 and N125. A, *right*) During remodeling, V95 interacts with P113 on the complementary subunit upon transitioning from closed to open pocket conformations. B) View as in A of the bacterial homolog sTeLIC, showing structures in the absence (PDB ID: 9EX6, red) and presence of 4-bromoamphetamine (BrAmp, PDB ID: 9EWL, blue). sTeLIC residues W75, Y104, and L93, corresponding to 5-HT_3A_R V95, N125, and P113 respectively, are shown. C, *left*) Frequency of contacts between 4-bromoamphetamine in site 1 and individual residues of the 5-HT_3A_R ECD in MD simulations. A contact is counted if two atoms of the ligand and receptor are within 4 °A of one another. C, *right*) Probability distribution of the minimum distance between the side-chain atoms of V95 and P113 on neighboring subunits, calculated over all stable MD simulation replicates associated with site 1. The distance captured in the cryo-EM structure (PDB ID: 6DG8) is highlighted red. D) Concentration-response curves for 5-HT activation of wild-type (WT) and mutant 5-HT_3A_Rs expressed in *Xenopus* oocytes. E) Percent modulation of WT, V95A, N125A, and P113G 5-HT_3_*_A_*R currents by 4-bromoamphetamine (n *≥* 5), relative to EC20 5-HT currents measured immediately before treatment. Asterisks indicate significance relative to WT (*P *≤* 0.05, **P *≤* 0.01, ***P *≤* 0.001, ****P *≤* 0.0001).

Local resolution is relatively poor in this region of *β*4 in multiple activated 5-HT_3A_R cryo-EM structures (Fig. S10), indicating it might be flexible and possibly sample such alternative conformations. Interestingly, this transition moved V95 roughly 4 ^°^A towards the subunit interface, bringing it into direct contact with residue P113 on *β*5 of the complementary subunit (Fig. 4A, C). A similar mechanism of side-chain rotation and intersubunit interaction was also captured in structures of sTeLIC (Fig. 4B), suggesting a generalizable model for pocket opening and ligand binding. Indeed, the inward-facing end of the Ω-loop was previously proposed to promote closed-to-open transitions of the 5-HT_3A_R by a similar mechanism, during which *β*4-F103 moves out of the vestibular site to interact with *β*6-P128 on the complementary subunit [33]. These effects are consistent with the general contraction of the upper ECD associated with pLGIC activation [2].

To functionally validate our putative ligand complex, we first engineered mutations at V95 and N125, two of the most frequent amino-acid contacts in simulations of this ligand pose (Fig. 4A,C). Substituting alanine at either of these positions weakened apparent serotonin sensitivity, increasing the agonist EC_50_ roughly 2-fold in 5-HT_3A_R expressing oocytes (Fig. 4D). Moreover, at serotonin concentrations giving equivalent levels of activation, either substitution significantly reduced potentiation by 600 nM–6 *µ*M 4-bromoamphetamine, rendering the receptor largely insensitive to the modulator (Fig. 4E). We further hypothesized that drug displacement of V95 towards P113 in the complementary subunit would promote interfacial contraction, thereby facilitating channel activation (Fig. 4A, C). In support of this model, substituting glycine at P113 dramatically suppressed potentiation by 4-bromoamphetamine, similar to its direct contacts in the vestibular site (Fig. 4E). Thus, functional recordings of site-directed mutants were in agreement with computational measurements of stability and accessibility in predicting a vestibular potentiating mechanism in the 5-HT_3A_R.

## Discussion

The resolution revolution has increasingly enabled single-particle cryo-EM reconstructions of previously inaccessible systems, including complex membrane proteins such as pLGICs [34]. While these advances have tremendous potential to launch a new era of structure-based drug design, relatively few structures have been determined in the presence of novel drugs. Visualizing drug binding at an atomic level, including the ligand and protein interactions and geometries, can support the design of pharmaceuticals that mimic the chemical structure, binding mode, and functional effect of known modulators. A persistent challenge in the field is that binding pockets may not be readily apparent in unliganded experimental structures, due to their transient nature combined with induced or selected fit binding mechanisms. Even when it is possible to resolve flexibility e.g. by cryo-EM, protein conformations compatible with binding might have too low probability to be detected. Therefore, to exploit the full potential of structural data, new types of methods are needed.

MD simulation and machine learning-based techniques offer notable opportunities to explore cryptic or otherwise experimentally obscure sites [18–21]. In MD simulations, a major challenge is the time scale required to transition a cryptic pocket from a closed to open state. Popular methods, in addition to FAST [22], include sampling water interfaces through scaled Hamiltonians (SWISH), in which nonbonded interactions of solvent molecules with protein atoms are progressively scaled, shifting the water properties toward more ligand-like behavior to promote cryptic-pocket opening [18]. A general challenge to MD-based methods is the need for computing resources and a system-specific simulation setup, particularly for larger protein complexes. To address this challenge, one recent study demonstrated the power of stochastic sub-sampling of multiple sequence alignment depth in AlphaFold2 combined with Markov state modeling to accelerate the discovery of novel cryptic pockets [20, 35]. Another study showed the effectiveness of training graph neural networks based on a large MD dataset to find cryptic pockets [19]. The FAST approach used here has been successfully applied to capturing pockets in Ebola viral protein 35 [36], SARS-CoV-2 [37] and the epidermal growth factor receptor [38], a notable diversity of targets.

Computational methods are inherently models offering predictions, and to validate these predictions we have used electrophysiology functional assays. This work demonstrates a pipeline that was able to reveal a functionally validated vestibular binding of an amphetamine derivative in a eukaryotic pLGIC. Amphetamines are widely used to treat diseases such as attention deficit hyperactivity disorder, narcolepsy, and irritable bowel syndrome [39]. However, several adverse side effects such as anorexia, weight loss, insomnia, and dependence have been reported [39]. The amphetamine-5-HT_3A_R complex proposed in this study could provide a starting point for designing analogous compounds to circumvent such side effects. Indeed, although its action has been largely attributed to metabotropic serotonin receptors, 4-bromoamphetamine produces toxicity in brainstem neurons [40] and depletes brain serotonin with a longer half-life than other amphetamines [41], supporting its relevance to mammalian neuropharmacology. Interestingly, a similar scaffold is featured in the contemporary recreational psychedelic 4-bromo-2,5-dimethoxyphenethylamine (2C-B) [42]. It remains to be seen whether other eukaryotic channels such as glycine, acetylcholine, or GABA_A_ receptors also contain modulatory sites in the vestibular region; application of our method to these systems could substantially enrich the development of pLGIC pharmacology.

Our results raise several mechanistic questions regarding vestibular modulation of 5-HT_3A_Rs. First, since we simulated pocket opening in the absence of a modulator, our study cannot conclusively distinguish whether drug binding occurs via a conformational selection, induced fit, or a combined mechanism. Metadynamics analysis could provide relevant insight, for instance by using our bound pose as a starting model and pocket opening/closing as well as drug binding/unbinding as reaction coordinates. Second, despite the fact that 4-bromoamphetamine alone is incapable of activating 5-HT_3A_R even at relatively high concentrations (Fig. 1C, Fig. S2), our study does not rule out the possibility that the ligand can also occupy the orthosteric serotonin-binding site. Further experimental studies may provide valuable insights on this matter.

Third, although our electrophysiology results showed that the ligand acts as a positive modulator, it remains unclear how ligand binding at the vestibular site may influence allosteric coupling between orthosteric serotonin binding and pore opening. Indeed, Since we were mainly interested in sampling the vestibular pocket in the ECD, located around 45 ^°^A from the TMD, we mildly restrained the M2 helix of the TMD to prevent commonly observed pore-collapsing events of pLGICs in MD simulations [43]. Dynamic allosteric coupling could in principle be investigated using MD-based methods such as Markov state modeling or metadynamics, with our putative complex as a starting point and employing polarizable or other forcefields that may better represent the hydration of the pore [44]. Given the evidence for cross-talk between the orthosteric and vestibular binding site in the bacterial channel GLIC [10–12], it would be interesting to see if a similar trend can be found in 5-HT_3A_R as well in the future. Overall, our results show how MD simulation-based methods can be used to sample previously unseen or cryptic binding pockets in cryo-EM structures and identify stable drug binding that can be validated with experimental assays. This appears to be a promising approach both for sampling binding pocket structural transitions for membrane proteins in general, and in particular to identify novel modulatory binding sites in the 5-HT_3A_R receptor and similar targets of high pharmaceutical relevance.

## Methods

### System preparations

Simulations were initiated from the serotonin-bound open-state cryo-EM structure of 5-HT_3_*_A_* (PDB ID 6DG8) [24]. The structure was embedded a symmetric membrane that approximates neuronal plasma membrane composition [45]: 44.4% cholesterol, 22.2% 1-palmitoyl-2-oleoyl-sn-glycero-3-phosphocholine (POPC), 22.2% 1-palmitoyl-2-oleoyl-sn-glycero-3-phosphoethanolamine (POPE), 10% 1-palmitoyl-2-oleoyl-sn-glycero-3-phospho-L-serine (POPS) and 1.1% phosphatidylinositol 4,5-bisphosphate (PtdIns(4,5)P2). The system was built using the Membrane Builder module of CHARMM-GUI [46] by solvating with TIP3P water [47] and neutralizing in 0.15 M NaCl to generate systems containing 300,000 atoms, with dimensions of 130 × 130 × 3 200 A . To allow for better sampling, 25 independent replicas were built by randomly configuring initial lipid placement around the protein using the Membrane mixer plugin in VMD [48, 49].

The systems were energy minimized and then relaxed in simulations at constant pressure (1 bar) and temperature (310K) for 30 ns, during which the position restraints on the protein and ligands were gradually released. The restraints were used as recommended by CHARMM-GUI. Then, production runs were performed with a mild position restraint of 50 kJ mol*^−^*^1^ nm*^−^*^2^ on the backbone atoms of the pore-facing residues at the M2 helix of the transmembrane domain (TMD), to prevent the commonly known pore-collapsing events of ligand-gated ion channels in MD simulations [43]. Also, a mild flat-bottom restraint of 20 kJ mol*^−^*^1^ nm*^−^*^2^ between the atoms of serotonin and residues at the binding sites was maintained, to prevent sudden release of GABA from the binding sites as been seen in our previous simulation.

### Adaptive sampling

To explore possible pocket openings at the vestibular site, we applied a goal-oriented adaptive sampling method, called fluctuation amplification of specific traits (FAST) [22]. Briefly, the method runs successive swarms of simulations where the starting points for each swarm are chosen from the set of all previously discovered conformations based on a reward function. This function balances (1) preferentially simulating structures with maximum pair-wise distances (Fig. 2A) to encourage the Ω-loop to adapt a more open conformation that may harbor cryptic pockets; with (2) a broad exploration of conformational space. The pair-wise distances were chosen from the residues located at the beta strands of ECD to the same at the Ω-loop, resulting in a total of 125 pairs, 25 per monomer. The broad exploration phase was implemented by favoring states that are poorly sampled compared to other states, based on the root root-mean-square deviation of the ECD residues. During FAST, we performed 30 generations of simulations with 25 simulations/generation and 40 ns/simulation, totaling 30 *µ*s. Since no biasing force is applied to any individual simulation, the final data set can be used to build a Markov state model (MSM) to extract the proper thermodynamics and kinetics [50–52], as detailed below.

### Markov state modeling

We used our trajectory dataset from FAST to construct a Markov state model (MSM) using pyEmma [53], by first featurizing the trajectory dataset using the 125 residue-residue distance pairs used for FAST sampling, as described above. The conformational space was then discretized into 1000 microstates using k-means clustering. Then, a transition probability matrix (TPM) was constructed by evaluating the probability of transitioning between each microstate within a lag time, *τ* . To choose an adequate lag time to construct a TPM that ensures Markovian behavior, multiple TPMs were first created using multiple maximum-likelihood MSMs with different lag times. The implied timescales were evaluated for each of these transition matrices, and saturation was observed at *τ* = 5 ns (Fig. S7). Thus, we built our final TPM using a maximum likelihood MSM with a lag time of 5 ns. This final TPM is symmetrized using a maximum likelihood approach to ensure detailed balance [53].

### MD simulations

MD simulations in this study were performed using GROMACS-2023 [54] utilizing CHARMM36m [26] and CHARMM36 [55] force field parameters for proteins and lipids, respectively. The force field parameters for the ligands were generated using the CHARMM General Force Field (CGenFF) [27–29]. Cation-*π* interaction-specific NBFIX parameters were used to maintain appropriate ligand-protein interactions at the aromatic cage, located at the binding sites [56]. Bonded and short-range non-bonded interactions were calculated every 2 fs, and periodic boundary conditions were employed in all three dimensions. The particle mesh Ewald (PME) method [57] was used to calculate long-range electrostatic interactions with a grid spacing below 0.1 nm*^−^*^3^. A force-based smoothing function was employed for pairwise nonbonded interactions at 1 nm with a cutoff of 1.2 nm. Pairs of atoms whose interactions were evaluated were searched and updated every 20 steps. A cutoff of 1.2 nm was applied to search for the interacting atom pairs. Constant pressure was maintained at 1 bar using the Parrinello-Rahman barostat [58] and temperature was kept at 300K with the v-rescale thermostat [59].

### Molecular docking

All the FAST sampled conformations were used for molecular docking of 4-bromoamphetamine using AutoDock Vina [60]. A grid box of dimensions of 24 x 22 _°_3 x 25 A at the vestibular site was employed for docking, outputting the best docking scored pose for each protein conformation.

### Analysis

System visualization and analysis were carried out using VMD and PyMOL [49, 61]. The “measure cluster” module implemented in VMD [62] was used for clustering analysis.

### Expression in oocytes and electrophysiology

The gene encoding mouse 5-HT_3_*_A_*R was inserted into the pBK-CMV expression vector. The desired mutation was created by site-directed mutagenesis using the Phusion High–Fidelity DNA Polymerase (ThermoFisher). The PCR product was digested overnight with DpnI at 37°C and transformed into XL1-Blue Supercompetent cells (Agilent) and the mutation was confirmed by sequencing (Eurofins Genomics). Oocytes from female Xenopus laevis frogs (Ecocyte Bioscience) were injected into the animal pole with 6 ng/32.2 nl DNA and incubated for 3–9 days at 13°C in MBS (88 mM NaCl, 1 mM KCl, 2.4 mM NaHCO_3_, 0.91 mM CaCl_2_, 0.82 mM MgSO_4_, 0.33 mM Ca(NO_3_)_2_, 10 mM HEPES, 0.5 mM theophylline, 0.1 mM G418, 17 mM streptomycin, 10,000 U/l penicillin and 2 mM sodium pyruvate, adjusted to pH 7.5) before two-electrode voltage-clamp (TEVC) recordings.

For TEVC recordings, glass electrodes filled with 3 M KCl (5 - 50 MΩ) were used to voltage-clamp the oocyte membrane potential at -70 mV with an OC-725C voltage clamp (Warner Instruments). Oocytes were continuously perfused with running buffer (123 mM NaCl, 2 mM KCl, and 2 mM MgSO_4_, and adjusted to pH 7.5) at a flow rate of 20 rpm. Currents were sampled and digitized at a sampling rate of 1 kHz with a Digidata 1440A. Current traces were plotted and analyzed by Clampfit (Molecular Devices).

For each BrAmp recording, a 1-M stock solution of BrAmp was prepared in dimethyl sulfoxide. The wash period between each 30-sec application was 10 min. After each BrAmp-containing application, the two subsequent applications were solely running buffer with *∼* EC20 5-HT to verify re-sensitization. To investigate whether BrAmp could directly activate 5-HT_3_*_A_*R, oocytes were washed with BrAmp for 1 min, then immediately exchanged to a *∼* EC20 5-HT-containing buffer with the same concentration of BrAmp for 30 s. Dose-response curves were fitted by nonlinear regression to the Boltzmann equation with variable slope and amplitudes using Prism 10.0.3 (GraphPad Software). Each reported value represents the mean *±* SEM for n *≥* four oocytes, and is analyzed with unpaired t-tests, with significant effects set at P *<* 0.05.

## Acknowledgments.

We thank Maxwell Zimmerman for his valuable feedback and discussion on the FAST simulation setup and troubleshooting. MD simulations were performed using computing facilities of the Karolina, LUMI and Discoverer Supercomputer through EuroHPC (grant nos. EHPC-REG-2023R01-103, EHPC-REG-2022R03-223 and EHPC-REG-2022R03-219, respectively) and the Swedish National Infrastructure for Computing (SNIC 2022/3-40) and supported by BioExcel (EuroHPC grant no. 101093290). N.H. was supported by a Marie Sklodowska-Curie Postdoctoral Fellowship (grant no. 101107036), E.K. by a Sven & Lily Lawski Foundation Doctoral Fellowship, and R.J.H., and E.L. by grants from the Swedish Research Council (grant nos. 2019-02433, 2021-05806) and Swedish e-Science Research Centre.

## Data and code availability

The raw MD simulation trajectories can be found on Zenodo: 10.5281/zenodo.10812994.

## Author contributions

N.H. designed research, performed the simulations and analyzed data; E.K. designed research, performed the experiments and analyzed data; and N.H., E.K., R.J.H., and E.L. wrote the paper. All authors (N.H., E.K., M.D., R.J.H., E.L.) read and contributed to finalizing the paper.

## Supplementary Figures and Tables

**Fig. S1.**
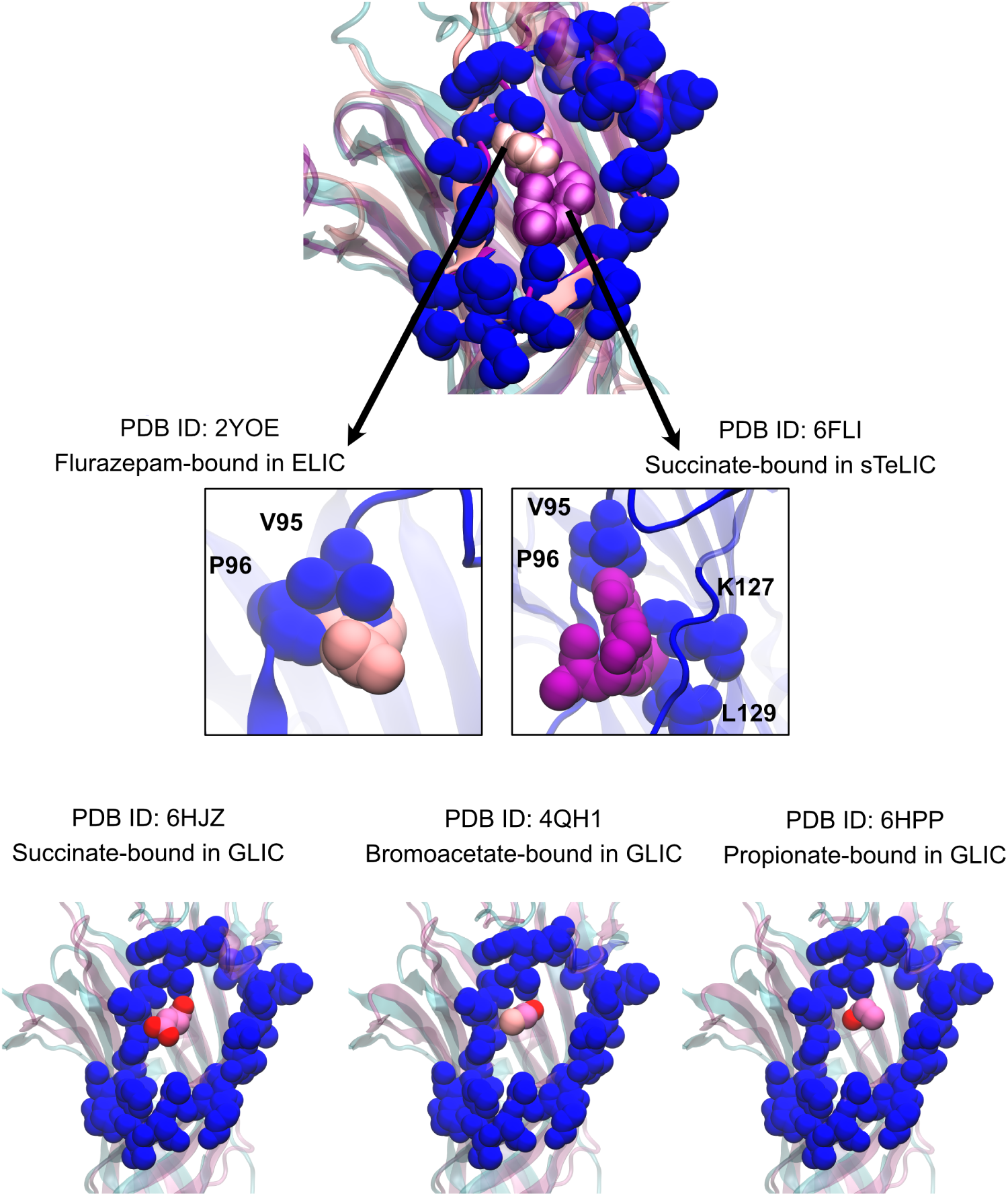
Structural alignment of putative vestibular sites in GLIC bound to different carboxylates (pink) and the 5-HT_3A_R (PDB ID: 6DG8, blue). In GLIC, spheres represent vestibule-bound ligands and in the 5-HT_3A_R, spheres represent amino-acid side chains in the Ω-loop.

**Fig. S2.**
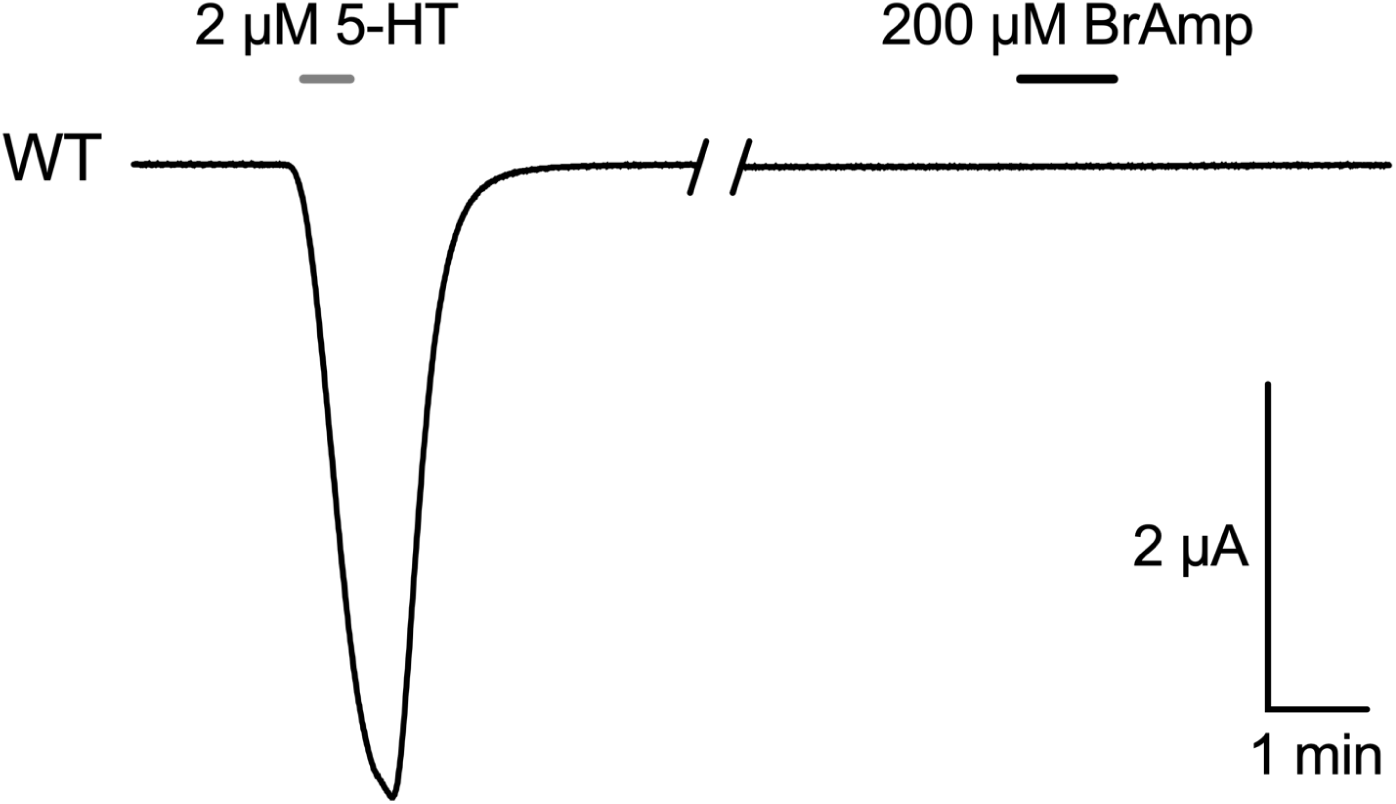
Sample oocyte electrophysiology trace showing a lack of direct activation by 4-bromoamphetamine up to 200 micromolar.

**Fig. S3.**
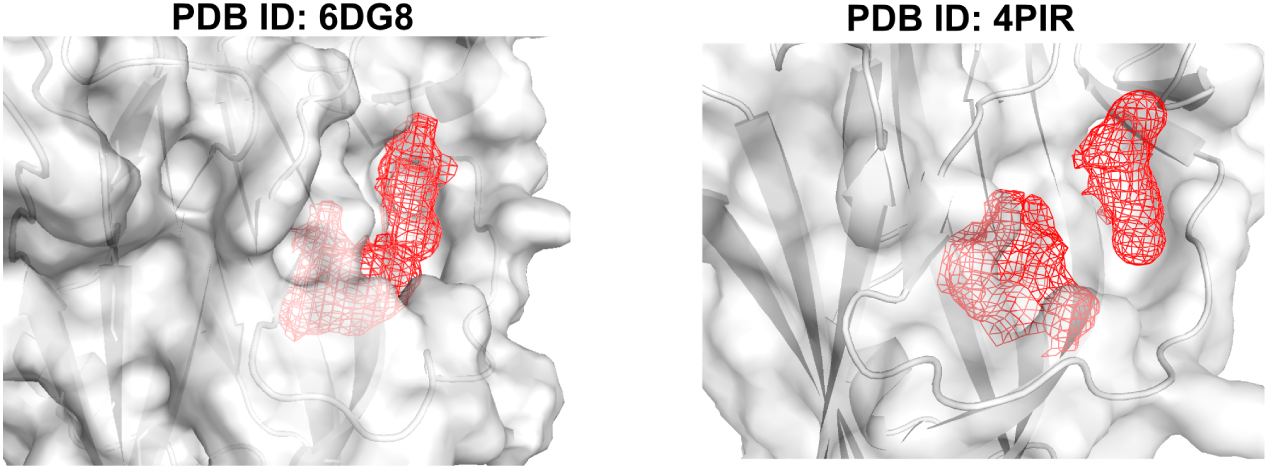
Pocket volumes at the 5-HT_3A_R vestibular site, generated in Fpocket [23], show no clear cavity for drug binding in two different activated experimental structures.

**Fig. S4.**
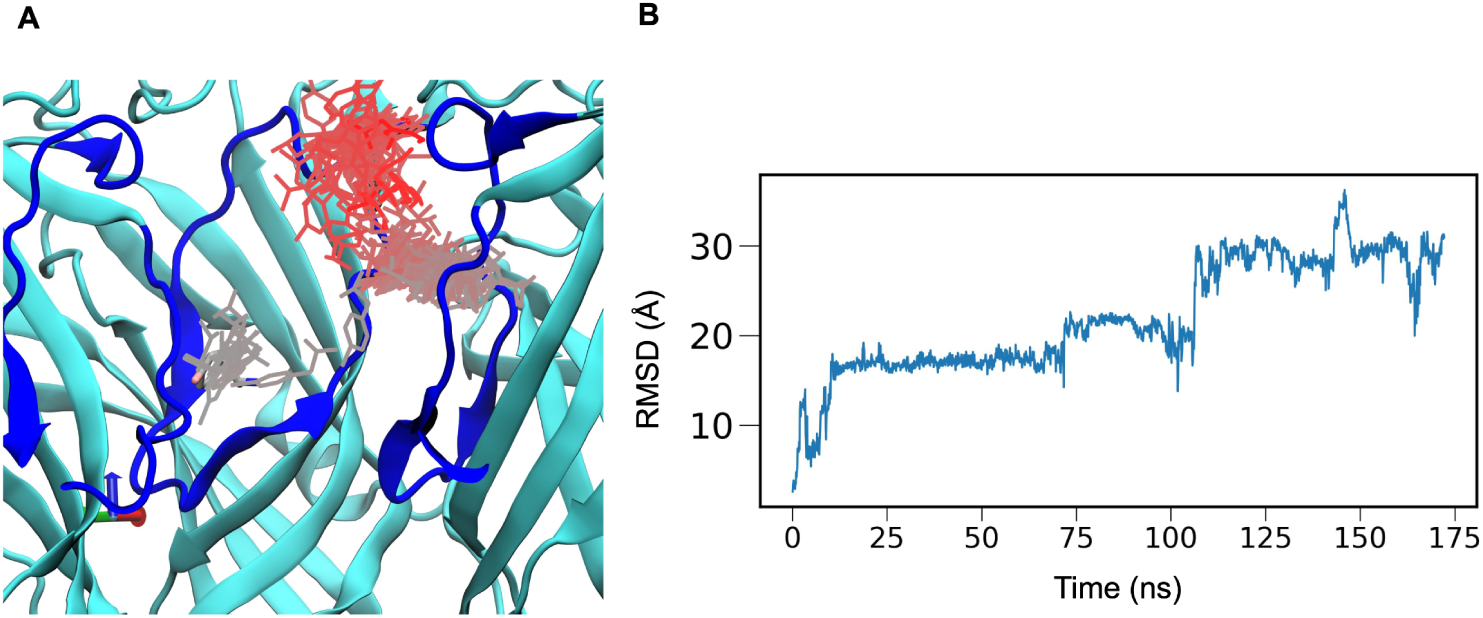
A) Docking of 4-bromoamphetamine to an activated-state 5-HT_3A_R (PDB ID: 6DG8) [24] did not produce stable binding, as illustrated by the time evolution of the ligand during simulation, colored by frame (white to red). B) RMSD of the ligand with respect to its original docked position during MD simulation.

**Fig. S5.**
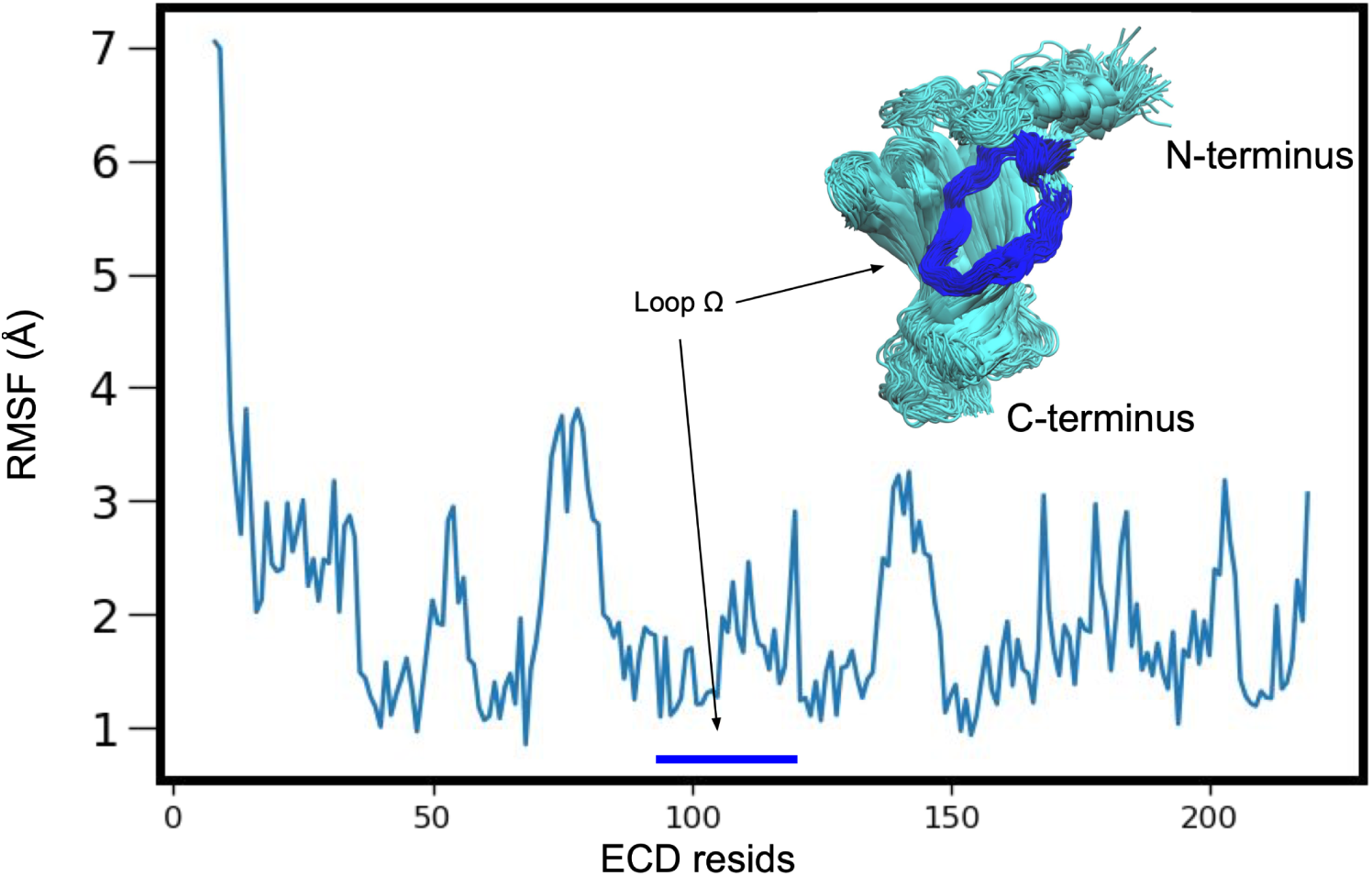
Root-mean-square fluctuation (RMSF) of ECD residues over the first generation of FAST simulations (25 replicates, 1 *µ*s total simulation time). Inset shows sampled ECD conformations as cyan ribbons, with the Ω-loop in blue.

**Fig. S6.**
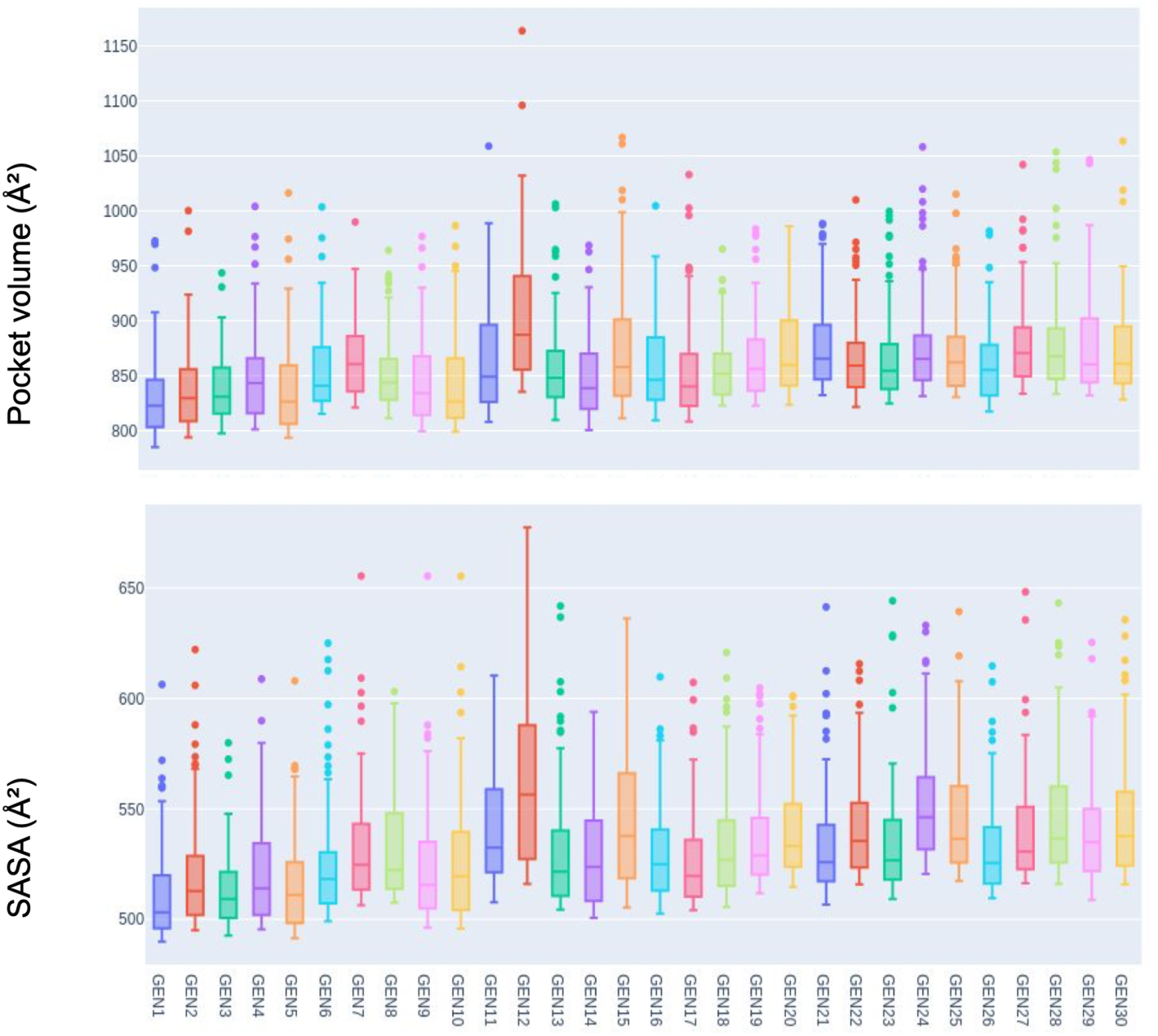
Box plot of pocket volumes and solvent-exposed surface areas in at the vestibular site, calculated using CAVER [32] for each FAST generation. Calculations were done with the highest 100 values for each generation.

**Fig. S7.**
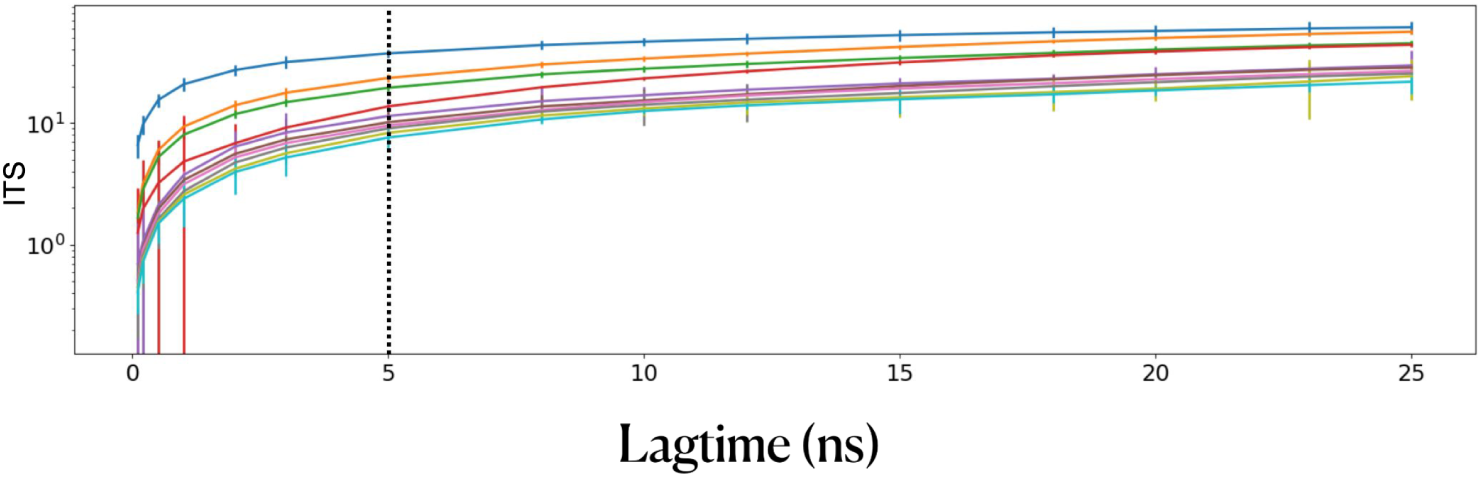
Implied timescales (ITS) plot of the top 10 slowest processes using multiple lag times in Markov state modeling of FAST sampling trajectories. Error bars indicate the uncertainty evaluated using a Bayesian estimated Markov state model. A lag time of 5 ns (dotted line) was chosen for Markov state model construction.

**Fig. S8.**
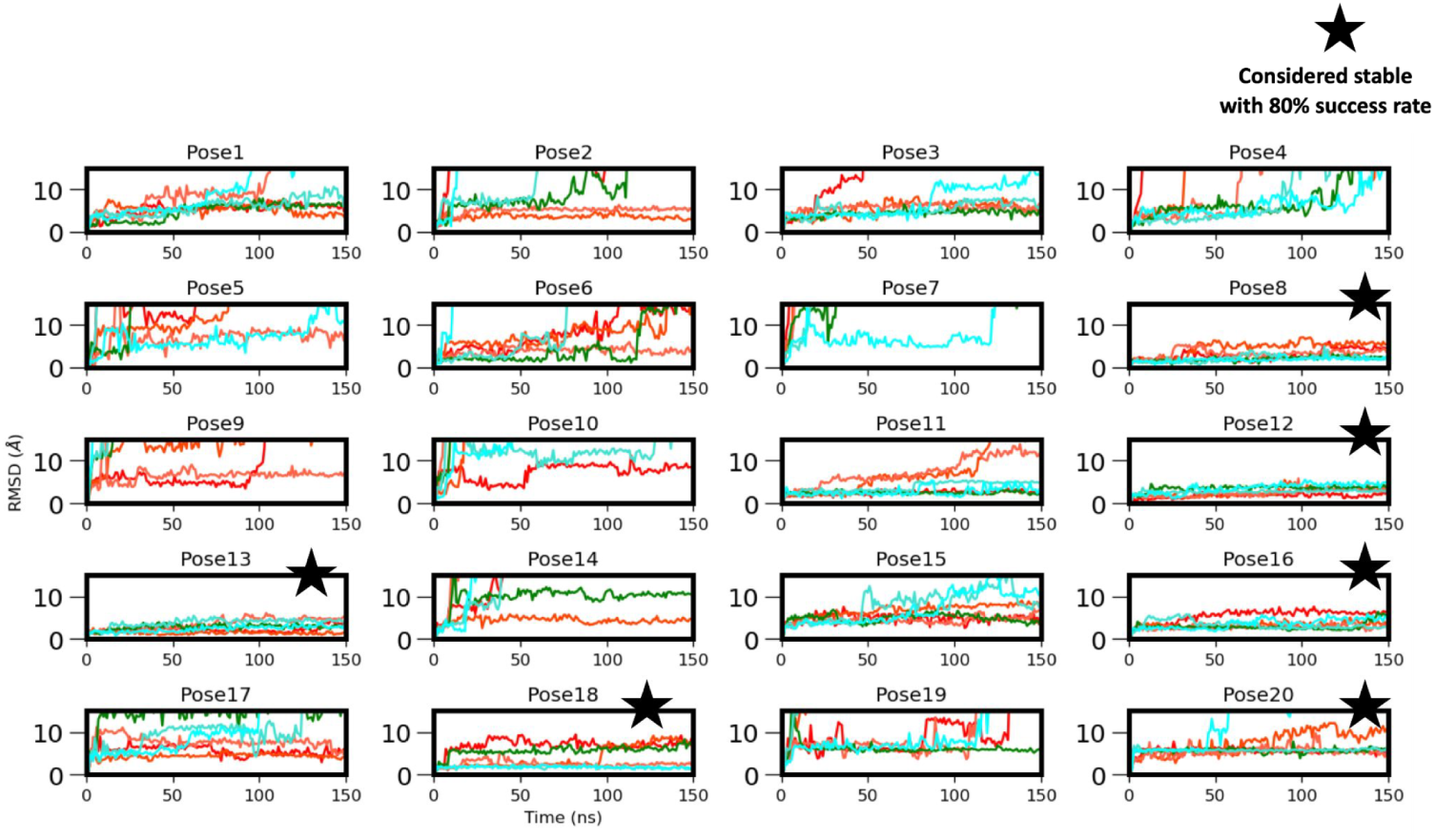
RMSD of 4-bromoamphetamine from its initial docked pose during MD simulations of each system simulated in three replicates each in CHARMM36 (shades of blue) and AMBER (shades of red). Stars indicate systems selected for further analysis due to remaining within 15 °A RMSD throughout at least 5 of 6 replicates,

**Fig. S9.**
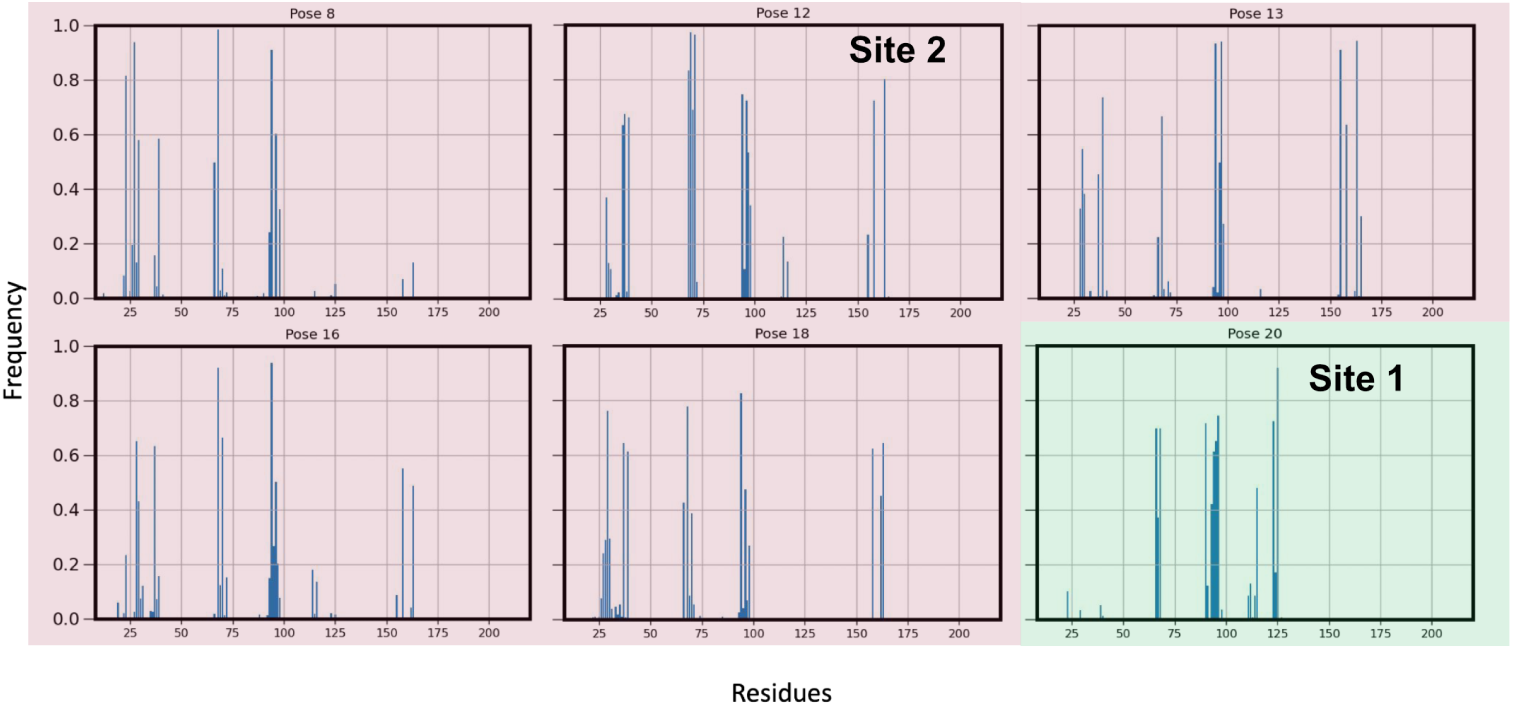
Frequency of contacts of 4-bromoamphetamine to individual residues of the 5-HT_3A_R in stable MD simulations of selected poses (Fig. S8). A contact is counted if two atoms of the ligand and receptor are within 4 °*A*. Green and red shading indicate categorization as sites 1 and 2, respectively, based on patterns of contacting residues.

**Fig. S10.**
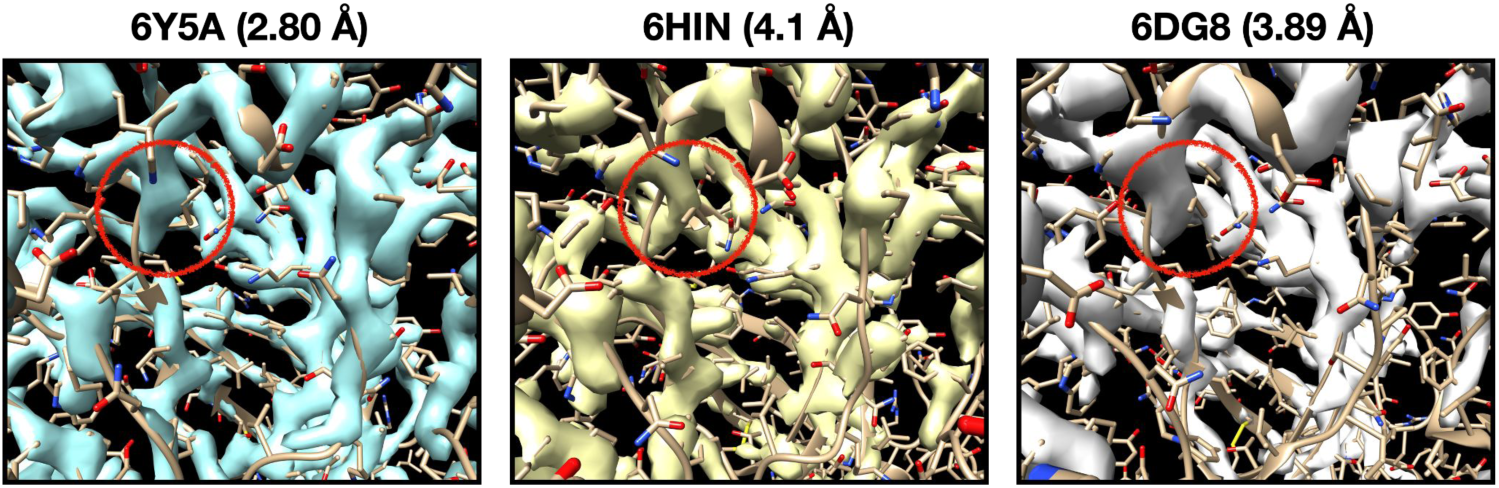
Cryo-EM densities and corresponding models for activated structures of the 5-HT_3A_R. Red circles highlight the densities around residue V95.

## References

[1] Lester, H.A., Dibas, M.I., Dahan, D.S., Leite, J.F., Dougherty, D.A.: Cys-loop receptors: new twists and turns. Trends in neurosciences 27(6), 329–336 (2004)

[2] Lynagh, T., Pless, S.A.: Principles of agonist recognition in Cys-loop receptors. Frontiers in physiology 5, 160 (2014)

[3] Zarkadas, E., Zhang, H., Cai, W., Effantin, G., Perot, J., Neyton, J., Chipot, C., Schoehn, G., Dehez, F., Nury, H.: The binding of palonosetron and other antiemetic drugs to the serotonin 5-HT3 receptor. Structure 28(10), 1131–1140 (2020)

[4] Sigel, E., Ernst, M.: The benzodiazepine binding sites of GABA*_A_* receptors. Trends in pharmacological sciences 39(7), 659–671 (2018)

[5] Sieghart, W., Savíc, M.M.: International union of basic and clinical pharmacology. CVI: GABAA receptor subtype-and function-selective ligands: key issues in translation to humans. Pharmacological reviews 70(4), 836–878 (2018)

[6] Kim, J.J., Gharpure, A., Teng, J., Zhuang, Y., Howard, R.J., Zhu, S., Noviello, C.M., Walsh Jr, R.M., Lindahl, E., Hibbs, R.E.: Shared structural mechanisms of general anaesthetics and benzodiazepines. Nature 585(7824), 303–308 (2020)

[7] Hu, H., Nemecz, Á, Van Renterghem, C., Fourati, Z., Sauguet, L., Corringer, P.-J., Delarue, M.: Crystal structures of a pentameric ion channel gated by alkaline pH show a widely open pore and identify a cavity for modulation. Proceedings of the National Academy of Sciences 115(17) (2018)

[8] Karlsson, E., Anden, O., Fan, C., Fourati, Z., Haouz, A., Zhuang, Y., Howard, R.J., Delarue, M., Lindahl, E.: Vestibular modulation by stimulant derivatives in a pentameric ligand-gated ion channel. bioRxiv, 592243 (2024). Preprint at 10.1101/2024.05.02.592243,

[9] Spurny, R., Ramerstorfer, J., Price, K., Brams, M., Ernst, M., Nury, H., Verheij, M., Legrand, P., Bertrand, D., Bertrand, S., Dougherty, D.A., Esch, I.J.P., Corringer, P.-J., Sieghart, W., Lummis, S.C.R., Ulens, C.: Pentameric ligand-gated ion channel ELIC is activated by GABA and modulated by benzodiazepines. Proceedings of the National Academy of Sciences 109(44), 3028–3034 (2012)

[10] Van Renterghem, C., Nemecz, Á., Delarue-Cochin, S., Joseph, D., Corringer, P.-J.: Fumarate as positive modulator of allosteric transitions in the pentameric ligand-gated ion channel GLIC: Requirement of an intact vestibular pocket. The Journal of physiology 601(12), 2447–2472 (2023)

[11] Fourati, Z., Sauguet, L., Delarue, M.: Structural evidence for the binding of monocarboxylates and dicarboxylates at pharmacologically relevant extracellular sites of a pentameric ligand-gated ion channel. Acta crystallographica section D: Structural Biology 76(7), 668–675 (2020)

[12] Renterghem, C.V., Nemecz, A., Madjebeur, K., Corringer, P.-J.: Short-chain mono-carboxylates as negative modulators of allosteric transitions in glic, and impact of a pre-*β*5 strand (loop *ω*) double mutation on crotonate, not butyrate effect. Physiological reports 12, 15916 (2024)

[13] Brams, M., Govaerts, C., Kambara, K., Price, K.L., Spurny, R., Gharpure, A., Pardon, E., Evans, G.L., Bertrand, D., Lummis, S.C., Hibbs, R.E., Steyaert, J., Ulens, C.: Modulation of the *Erwinia* ligand-gated ion channel (ELIC) and the 5-HT_3_ receptor via a common vestibule site. eLife 9, 51511 (2020)

[14] Morales-Perez, C.L., Noviello, C.M., Hibbs, R.E.: X-ray structure of the human *α*4*β*2 nicotinic receptor. Nature 538(7625), 411–415 (2016)

[15] Walsh Jr, R.M., Roh, S.-H., Gharpure, A., Morales-Perez, C.L., Teng, J., Hibbs, R.E.: Structural principles of distinct assemblies of the human α4β2 nicotinic receptor. Nature 557(7704), 261–265 (2018)

[16] Du, J., Lü, W., Wu, S., Cheng, Y., Gouaux, E.: Glycine receptor mechanism elucidated by electron cryo-microscopy. Nature 526(7572), 224–229 (2015)

[17] Hassaine, G., Deluz, C., Grasso, L., Wyss, R., Tol, M.B., Hovius, R., Graff, A., Stahlberg, H., Tomizaki, T., Desmyter, A., Moreau, C., Li, X.-D., Poitevin, F., Vogel, H., Nury, H.: X-ray structure of the mouse serotonin 5-HT_3_ receptor. Nature 512(7514), 276–281 (2014)

[18] Oleinikovas, V., Saladino, G., Cossins, B.P., Gervasio, F.L.: Understanding cryptic pocket formation in protein targets by enhanced sampling simulations. Journal of the American Chemical Society 138(43), 14257–14263 (2016)

[19] Meller, A., Ward, M.D., Borowsky, J.H., Lotthammer, J.M., Kshirsagar, M., Oviedo, F., Ferres, J.L., Bowman, G.: Predicting the locations of cryptic pockets from single protein structures using the PocketMiner graph neural network. Biophysical journal 122(3), 445 (2023)

[20] Meller, A., Bhakat, S., Solieva, S., Bowman, G.R.: Accelerating cryptic pocket discovery using AlphaFold. Journal of chemical theory and computation 19(14), 4355–4363 (2023)

[21] Kuzmanic, A., Bowman, G.R., Juarez-Jimenez, J., Michel, J., Gervasio, F.L.: Investigating cryptic binding sites by molecular dynamics simulations. Accounts of chemical research 53(3), 654–661 (2020)

[22] Zimmerman, M.I., Bowman, G.R.: FAST conformational searches by balancing exploration/exploitation trade-offs. Journal of chemical theory and computation 11(12), 5747–5757 (2015)

[23] Le Guilloux, V., Schmidtke, P., Tuffery, P.: Fpocket: an open source platform for ligand pocket detection. BMC bioinformatics 10(1), 1–11 (2009)

[24] Basak, S., Gicheru, Y., Rao, S., Sansom, M.S., Chakrapani, S.: Cryo-EM reveals two distinct serotonin-bound conformations of full-length 5-HT_3*A*_ receptor. Nature 563(7730), 270–274 (2018)

[25] Hart, K.M., Ho, C.M., Dutta, S., Gross, M.L., Bowman, G.R.: Modelling proteins’ hidden conformations to predict antibiotic resistance. Nature communications 7(1), 12965 (2016)

[26] Huang, J., Rauscher, S., Nawrocki, G., Ran, T., Feig, M., De Groot, B.L., Grubmüller, H., MacKerell Jr, A.D.: CHARMM36m: an improved force field for folded and intrinsically disordered proteins. Nature methods 14(1), 71–73 (2017)

[27] Vanommeslaeghe, K., Raman, E.P., MacKerell Jr, A.D.: Automation of the CHARMM general force field (CGenFF) II: assignment of bonded parameters and partial atomic charges. Journal of chemical information and modeling 52(12), 3155–3168 (2012)

[28] Vanommeslaeghe, K., Hatcher, E., Acharya, C., Kundu, S., Zhong, S., Shim, J., Darian, E., Guvench, O., Lopes, P., Vorobyov, I., Mackerell Jr, A.D.: CHARMM general force field: A force field for drug-like molecules compatible with the CHARMM all-atom additive biological force fields. Journal of computational chemistry 31(4), 671–690 (2010)

[29] Vanommeslaeghe, K., MacKerell Jr, A.D.: Automation of the CHARMM general force field (CGenFF) I: bond perception and atom typing. Journal of chemical information and modeling 52(12), 3144–3154 (2012)

[30] Tian, C., Kasavajhala, K., Belfon, K.A., Raguette, L., Huang, H., Migues, A.N., Bickel, J., Wang, Y., Pincay, J., Wu, Q., Simmerling, C.: ff19SB: Amino-acid-specific protein backbone parameters trained against quantum mechanics energy surfaces in solution. Journal of chemical theory and computation 16(1), 528–552 (2019)

[31] He, X., Man, V.H., Yang, W., Lee, T.-S., Wang, J.: A fast and high-quality charge model for the next generation general AMBER force field. The Journal of chemical physics 153(11) (2020)

[32] Chovancova, E., Pavelka, A., Benes, P., Strnad, O., Brezovsky, J., Kozlikova, B., Gora, A., Sustr, V., Klvana, M., Medek, P., Biedermannova, L., Sochor, J., Damborsky, J.: CAVER 3.0: a tool for the analysis of transport pathways in dynamic protein structures. PLoS computational biology 8(10), 1002708 (2012)

[33] Polovinkin, L., Hassaine, G., Perot, J., Neumann, E., Jensen, A.A., Lefebvre, S.N., Corringer, P.-J., Neyton, J., Chipot, C., Dehez, F., Schoehn, G., Nury, H.: Conformational transitions of the serotonin 5-HT_3_ receptor. Nature 563(7730), 275–279 (2018)

[34] Howard, R.J.: Elephants in the dark: Insights and incongruities in pentameric ligand-gated ion channel models. Journal of Molecular Biology 433(17), 167128 (2021)

[35] Jumper, J., Evans, R., Pritzel, A., Green, T., Figurnov, M., Ronneberger, O., Tunyasuvunakool, K., Bates, R., Žídek, A., Potapenko, A., Bridgland, A., Meyer, C., Kohl, S.A.A., Ballard, A.J., Cowie, A., Romera-Paredes, B., Nikolov, S., Jain, R., Adler, J., Back, T., Petersen, S., Reiman, D., Clancy, E., Zielinski, M., Steinegger, M., Pacholska, M., Berghammer, T., Bodenstein, S., Silver, D., Vinyals, O., Senior, A.W., Kavukcuoglu, K., Kohli, P., Hassabis, D.: Highly accurate protein structure prediction with AlphaFold. Nature 596(7873), 583–589 (2021)

[36] Cruz, M.A., Frederick, T.E., Mallimadugula, U.L., Singh, S., Vithani, N., Zimmerman, M.I., Porter, J.R., Moeder, K.E., Amarasinghe, G.K., Bowman, G.R.: A cryptic pocket in ebola VP35 allosterically controls RNA binding. Nature Communications 13(2269) (2022)

[37] Zimmerman, M., Porter, J., Ward, M., Singh, S., Vithani, N., Meller, A., Mallimadugula, U., Kuhn, C., Borowsky, J., Wiewiora, R., Hurley, M., Harbison, A., Fogarty, C., Coffland, J., Fadda, E., Voelz, V., Chodera, J., Bowman, G.: SARS-CoV-2 simulations go exascale to predict dramatic spike opening and cryptic pockets across the proteome. Nature Chemistry 13, 651–659 (2021)

[38] Behring, J.B., Post, S., Mooradian, A.D., Egan, M.J., Zimmerman, M.I., Clements, J.L., Bowman, G.R., Held, J.M.: Spatial and temporal alterations in protein structure by EGF regulate cryptic cysteine oxidation. Science signaling 13(615), 7315 (2020)

[39] Heal, D.J., Smith, S.L., Gosden, J., Nutt, D.J.: Amphetamine, past and present – a pharmacological and clinical perspective. Journal of psychopharmacology 27(6), 479–496 (2013)

[40] Harvey, J.A., McMaster, S.E., Fuller, R.W.: Comparison between the neurotoxic and serotonin-depleting effects of various halogenated derivatives of amphetamine in the rat. Journal of pharmacology and experimental therapeutics 202(3), 581– 589 (1977)

[41] Fuller, R.W., Baker, J.C., Perry, K.W., Molloy, B.B.: Comparison of 4-chloro-, 4-bromo- and 4-fluoroamphetamine in rats: drug levels in brain and effects on brain serotonin metabolism. Neuropharmacology 14(10), 739–746 (1975)

[42] Dean, B.V., Stellpflug, S.J., Burnett, A.M., Engebretsen, K.M.: 2c or not 2c: phenethylamine designer drug review. Journal of Medical Toxicology 9, 172–178 (2013)

[43] Dämgen, M.A., Biggin, P.C.: A refined open state of the glycine receptor obtained via molecular dynamics simulations. Structure 28(1), 130–139 (2020)

44. Jing, Z., Liu, C., Cheng, S.Y., Qi, R., Walker, B.D., Piquemal, J.-P., Ren, P.: Polarizable force fields for biomolecular simulations: Recent advances and applications. Annual Review of biophysics 48, 371–394 (2019)

[45] Inǵolfsson, H.I., Carpenter, T.S., Bhatia, H., Bremer, P.-T., Marrink, S.J., Lightstone, F.C.: Computational lipidomics of the neuronal plasma membrane. Biophysical journal 113(10), 2271–2280 (2017)

[46] Jo, S., Kim, T., Iyer, V.G., Im, W.: CHARMM-GUI: a web-based graphical user interface for CHARMM. Journal of computational chemistry 29(11), 1859–1865 (2008)

[47] Jorgensen, W.L., Chandrasekhar, J., Madura, J.D., Impey, R.W., Klein, M.L.: Comparison of simple potential functions for simulating liquid water. The Journal of chemical physics 79(2), 926–935 (1983)

[48] Licari, G., Dehghani-Ghahnaviyeh, S., Tajkhorshid, E.: Membrane mixer: a toolkit for efficient shuffling of lipids in heterogeneous biological membranes. Journal of Chemical Information and Modeling 62(4), 986–996 (2022)

[49] Humphrey, W., Dalke, A., Schulten, K.: VMD: visual molecular dynamics. Journal of molecular graphics 14(1), 33–38 (1996)

[50] Huang, X., Bowman, G.R., Bacallado, S., Pande, V.S.: Rapid equilibrium sampling initiated from nonequilibrium data. Proceedings of the National Academy of Sciences 106(47), 19765–19769 (2009)

[51] Bowman, G.R., Pande, V.S., Nóe, F.: An introduction to markov state models and their application to long timescale molecular simulation. Springer Science & Business Media 797 (2013)

[52] Nóe, F., Schütte, C., Vanden-Eijnden, E., Reich, L., Weikl, T.R.: Constructing the equilibrium ensemble of folding pathways from short off-equilibrium simulations. Proceedings of the National Academy of Sciences 106(45), 19011–19016 (2009)

[53] Scherer, M.K., Trendelkamp-Schroer, B., Paul, F., Pérez-Herńandez, G., Hoffmann, M., Plattner, N., Wehmeyer, C., Prinz, J.-H., Nóe, F.: PyEMMA 2: A software package for estimation, validation, and analysis of Markov models. Journal of chemical theory and computation 11(11), 5525–5542 (2015)

[54] Páll, S., Zhmurov, A., Bauer, P., Abraham, M., Lundborg, M., Gray, A., Hess, B., Lindahl, E.: Heterogeneous parallelization and acceleration of molecular dynamics simulations in GROMACS. The Journal of chemical physics 153(13), 134110 (2020)

[55] Klauda, J.B., Venable, R.M., Freites, J.A., O’Connor, J.W., Tobias, D.J., Mondragon-Ramirez, C., Vorobyov, I., MacKerell Jr, A.D., Pastor, R.W.: Update of the CHARMM all-atom additive force field for lipids: validation on six lipid types. The journal of physical chemistry B 114(23), 7830–7843 (2010)

[56] Liu, H., Fu, H., Chipot, C., Shao, X., Cai, W.: Accuracy of alternate nonpolarizable force fields for the determination of protein-ligand binding affinities dominated by cation-*π* interactions. Journal of chemical theory and computation 17(7), 3908–3915 (2021)

[57] Darden, T., York, D., Pedersen, L.: Particle mesh Ewald: An N ·log(N) method for Ewald sums in large systems. The Journal of chemical physics 98(12), 10089–10092 (1993)

[58] Parrinello, M., Rahman, A.: Crystal structure and pair potentials: A molecular-dynamics study. Physical review letters 45(14), 1196 (1980)

[59] Bussi, G., Donadio, D., Parrinello, M.: Canonical sampling through velocity rescaling. The Journal of chemical physics 126(1), 014101 (2007)

[60] Trott, O., Olson, A.J.: Autodock Vina: Improving the speed and accuracy of docking with a new scoring function, efficient optimization, and multithreading. Journal of computational chemistry 31(2), 455–461 (2010)

[61] DeLano, W.L.: PyMOL: An open-source molecular graphics tool. CCP4 Newsl. Protein Crystallogr 40(1), 82–92 (2002)

[62] Heyer, L.J., Kruglyak, S., Yooseph, S.: Exploring expression data: identification and analysis of coexpressed genes. Genome research 9(11), 1106–1115 (1999)

