## Supplementary data for "Discovering cryptic pocket opening and binding of a stimulant derivative in a vestibular site of the 5-HT_3_*_A_* receptor"

### Supplementary Figures and Tables

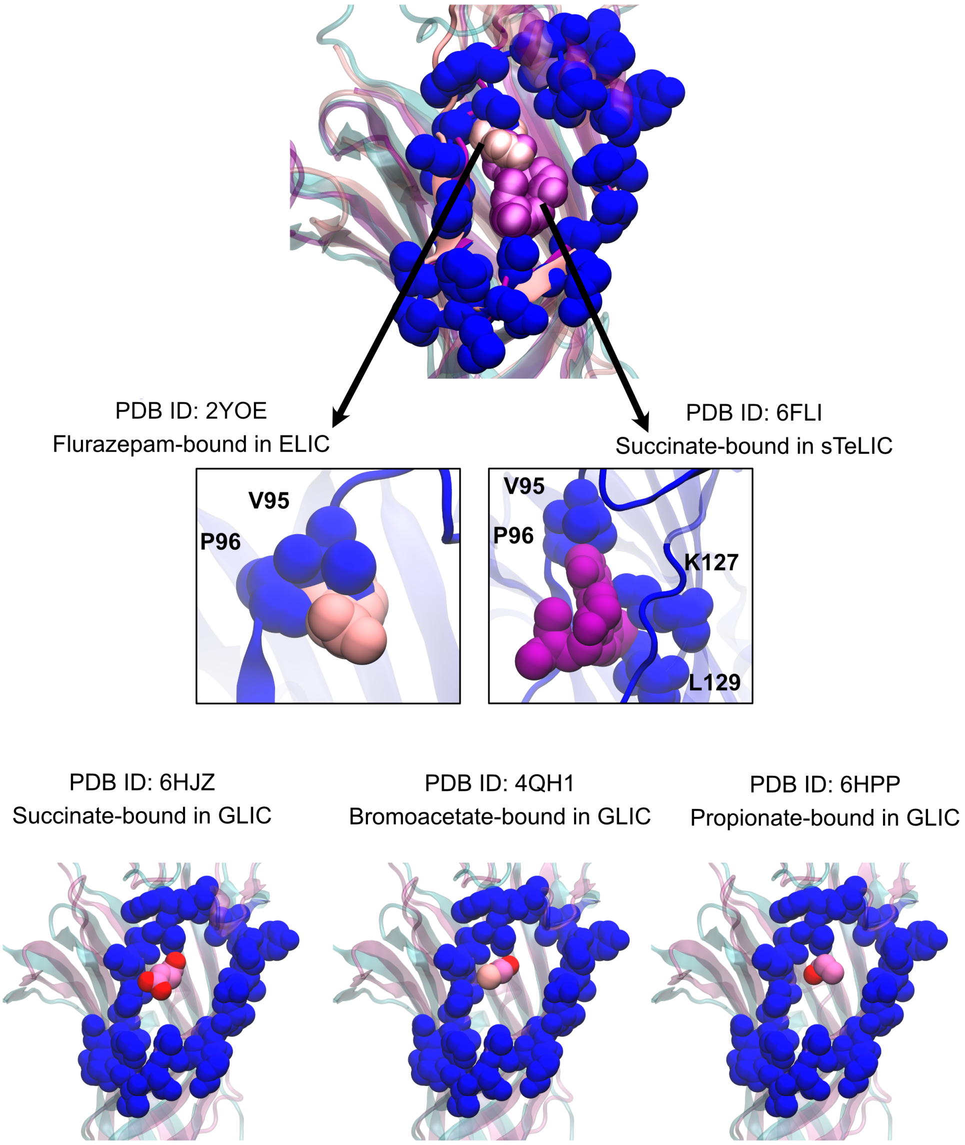


**Fig. S1** Structural alignment of putative vestibular sites in GLIC bound to different carboxylates (pink) and the 5-HT$_{3A}$R (PDB ID: 6DG8, blue). In GLIC, spheres represent vestibule-bound ligands and in the 5-HT$_{3A}$R, spheres represent amino-acid side chains in the Ω-loop.

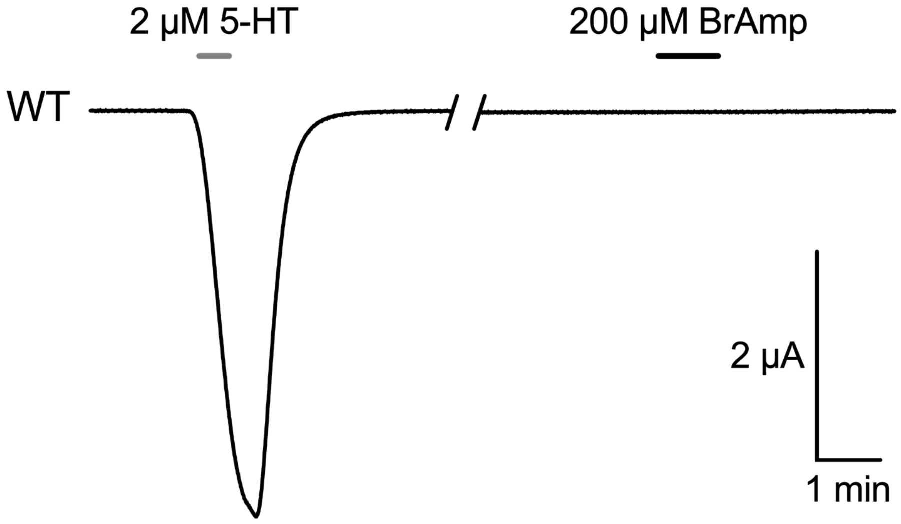


**Fig. S2** Sample oocyte electrophysiology trace showing a lack of direct activation by 4-bromoamphetamine up to 200 micromolar.

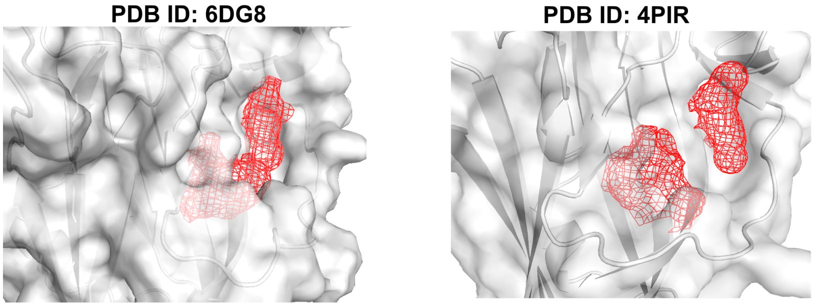


**Fig. S3** Pocket volumes at the 5-$HT_{3A}R$ vestibular site, generated in Fpocket [23], show no clear cavity for drug binding in two different activated experimental structures.

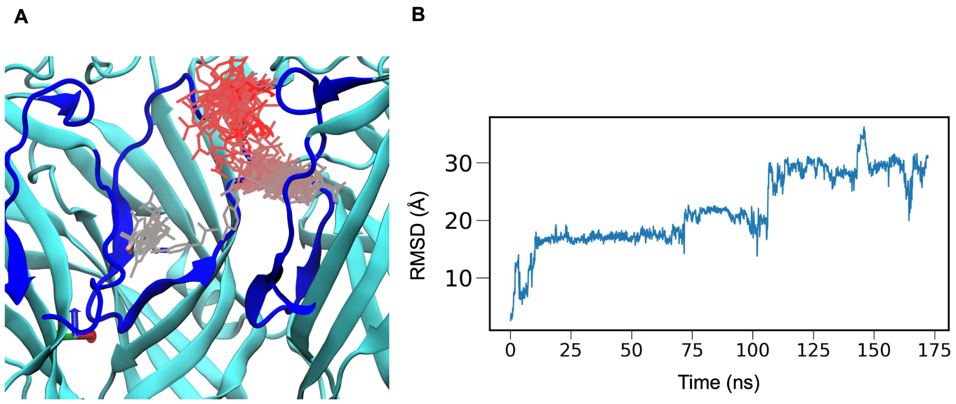


**Fig. S4** A) Docking of 4-bromoamphetamine to an activated-state 5-$HT_{3A}$R (PDB ID: 6DG8) [24] did not produce stable binding, as illustrated by the time evolution of the ligand during simulation, colored by frame (white to red). B) RMSD of the ligand with respect to its original docked position during MD simulation.

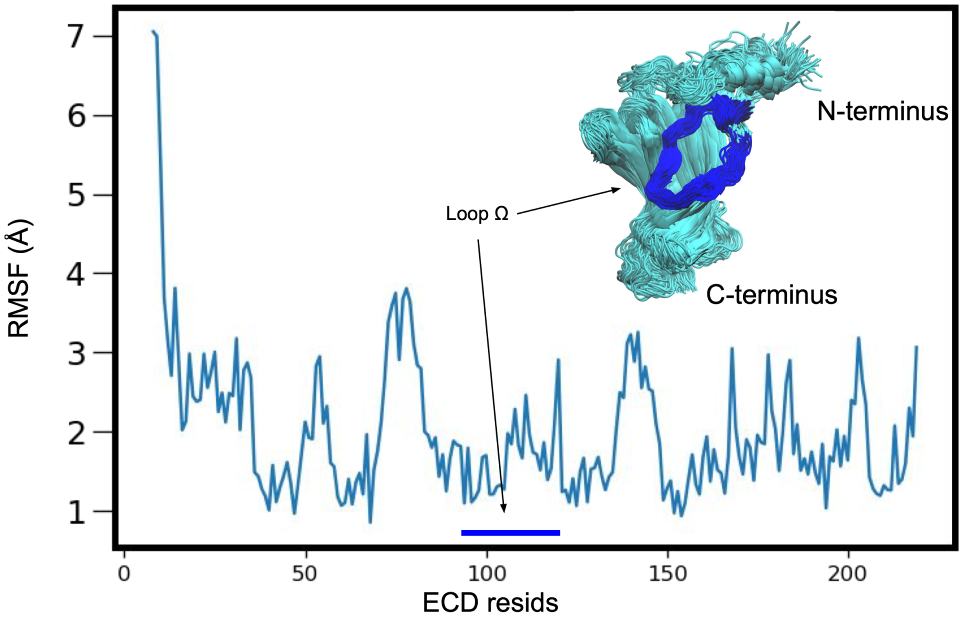


**Fig. S5** Root-mean-square fluctuation (RMSF) of ECD residues over the first generation of FAST simulations (25 replicates, 1 $\mu$s total simulation time). Inset shows sampled ECD conformations as cyan ribbons, with the $\Omega$-loop in blue.

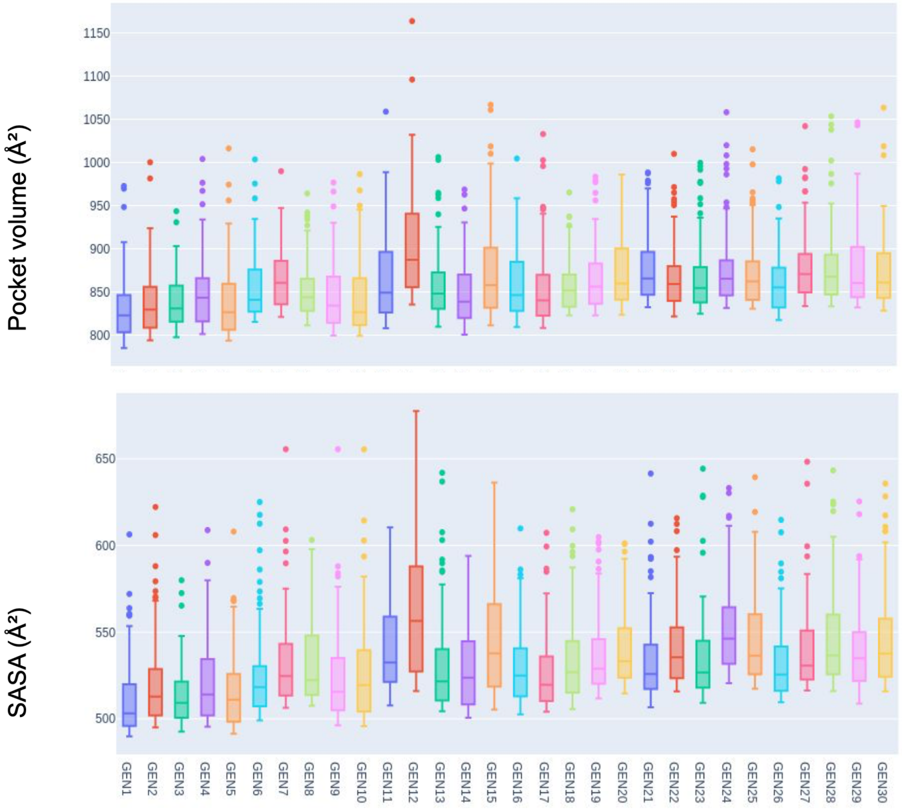


**Fig. S6** Box plot of pocket volumes and solvent-exposed surface areas in at the vestibular site, calculated using CAVER [32] for each FAST generation. Calculations were done with the highest 100 values for each generation.

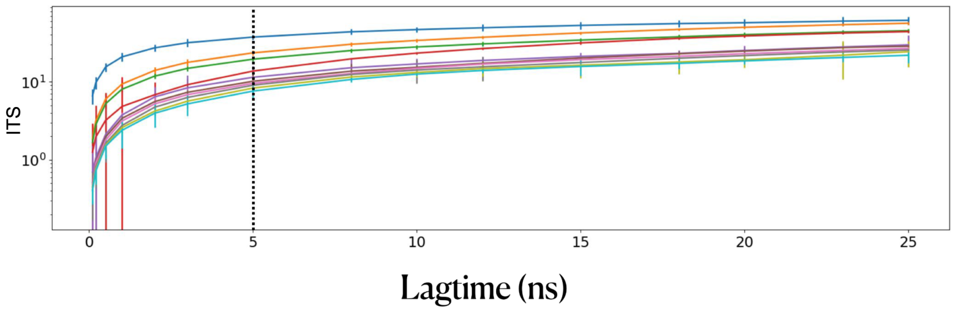


**Fig. S7** Implied timescales (ITS) plot of the top 10 slowest processes using multiple lag times in Markov state modeling of FAST sampling trajectories. Error bars indicate the uncertainty evaluated using a Bayesian estimated Markov state model. A lag time of 5 ns (dotted line) was chosen for Markov state model construction.

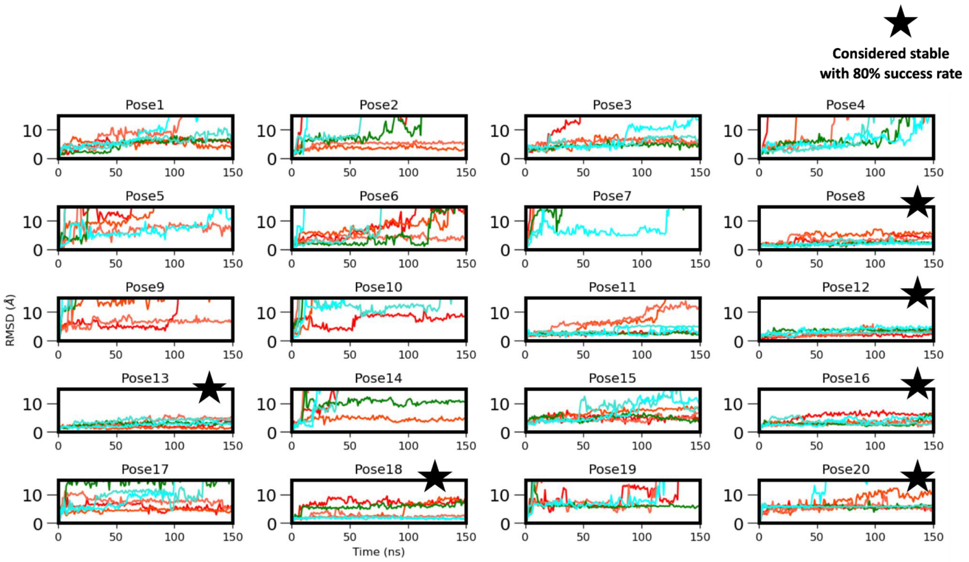


**Fig. S8** RMSD of 4-bromoamphetamine from its initial docked pose during MD simulations of each system simulated in three replicates each in CHARMM36 (shades of blue) and AMBER (shades of red). Stars indicate systems selected for further analysis due to remaining within 15 Å RMSD throughout at least 5 of 6 replicates,

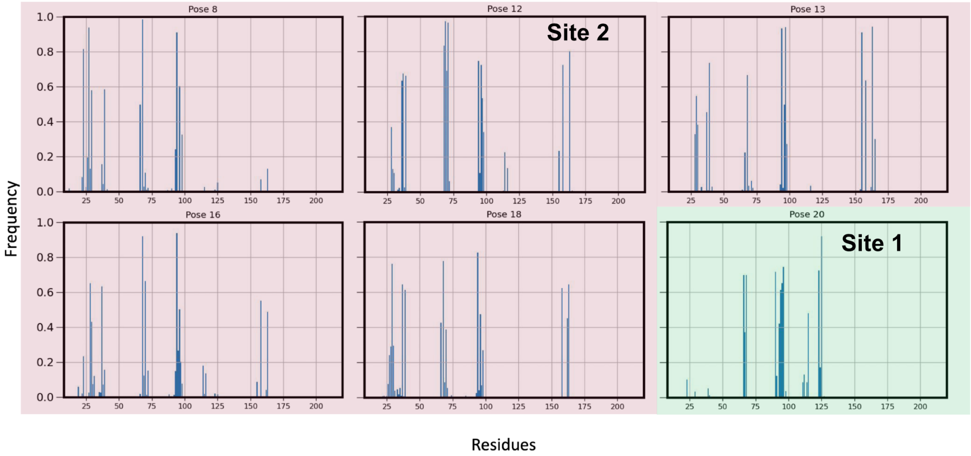


**Fig. S9** Frequency of contacts of 4-bromoamphetamine to individual residues of the 5-$HT_{3A}R$ in stable MD simulations of selected poses (Fig. S8). A contact is counted if two atoms of the ligand and receptor are within 4 $\AA$. Green and red shading indicate categorization as sites 1 and 2, respectively, based on patterns of contacting residues.

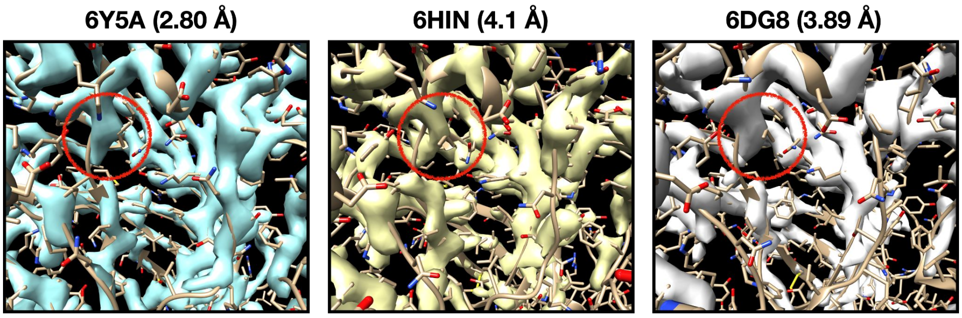


**Fig. S10** Cryo-EM densities and corresponding models for activated structures of the 5-$HT_{3A}$R. Red circles highlight the densities around residue V95.
